# Tau pathology in the dorsal raphe may be a prodromal indicator of Alzheimer’s disease

**DOI:** 10.1101/2022.11.22.517403

**Authors:** Samantha Pierson, Kimberly L. Fiock, Ruixiang Wang, Nagalakshmi Balasubramanian, Jessica Reindhardt, Kanza M. Khan, Thomas D. James, Ryan Betters, Kaancan Deniz, Gloria Lee, Georgina Aldridge, Marco M. Hefti, Catherine A. Marcinkiewcz

## Abstract

Protein aggregation in brainstem nuclei is thought to occur in the early stages of Alzheimer’s disease (AD), but its specific role in driving prodromal symptoms and disease progression is largely unknown. The dorsal raphe nucleus (DRN) contains a large population of serotonin (5-hydroxytryptamine; 5-HT) neurons that regulate mood, reward-related behavior, and sleep, which are all disrupted in AD. We report here that tau pathology is present in the DRN of individuals 54-80 years old without a known history of dementia and was found at higher frequency than α-synuclein and TDP-43. Most AD cases had tau pathology in the DRN (90%), whereas only a subset contained TDP-43 or α-synuclein, but not both (30%). To evaluate how early tau pathology impacts behavior, we overexpressed human P301L-tau in the DRN of mice and observed depressive-like behaviors and hyperactivity without any deficits in spatial memory. Tau pathology was predominantly found in neurons relative to glia and colocalized with a significant proportion of Tph2-expressing neurons in the DRN. 5-HT neurons were also hyperexcitable in P301L-tau^DRN^ mice, and there was an increase in the amplitude of excitatory post-synaptic currents (EPSCs), suggestive of increased glutamatergic transmission. Moreover, astrocytic density was elevated in the DRN and accompanied by an increase in IL-1α and Frk expression, which is indicative of inflammation. Additionally, tau pathology was detected in axonal processes in the thalamus, hypothalamus, amygdala, and caudate putamen and a significant proportion of this tau pathology colocalized with the serotonin reuptake transporter (SERT), suggesting that tau may spread in an anterograde manner to regions outside the DRN. Together these results indicate that tau pathology accumulates in the DRN in a subset of individuals over 50 years and may lead to behavioral dysregulation, 5-HT neuronal dysfunction, and activation of local astrocytes which may be prodromal indicators of AD.

## Introduction

Alzheimer’s disease (AD) is an age-related neurodegenerative disease marked by memory loss and cognitive decline[1]. Neuropathological changes in the brain occur before any notable deficits in memory and cognition, making clinical diagnosis during the early stages of AD a significant challenge. New disease-modifying therapies show promise but require early administration, so identifying biomarkers of prodromal AD that would enable earlier diagnosis is one of the highest priorities in the field. Neuropsychiatric symptoms (NPS) have been reported in the early stages of AD and may be a useful indicator of underlying neuropathology and imminent cognitive decline[2–6]. While the neural circuits underlying NPS are currently unknown, several lines of evidence suggest that the dorsal raphe nucleus (DRN) may be critically involved. The DRN is a brainstem nucleus with extensive efferent serotonergic projections to the forebrain that have been shown to regulate mood, anxiety, and reward-related behavior[7–11]. Neurofibrillary tangles (NFTs) are present in the DRN at an early stage (Braak Stage 0) before they appear in the entorhinal cortex and other structures [12], and at later Braak stages there is marked loss of serotonin (5-hydroxytryptamine; 5-HT) neurons, serotonin transporter (SERT) binding sites, and 5-HT_1A_ receptor binding sites [13–15]. Still, the presence of DRN NFTs in early AD remains underreported in the literature, and the timing of its appearance and prevalence in the general population is unknown. Another outstanding question is whether DRN tau pathology can alter cognitive function or induce behavioral symptoms associated with prodromal AD. In the present study, we examined post-mortem DRN sections for tau, synuclein, and TDP-43 pathology in a group of individuals aged 54-80 years with no documented history of neurological disease as well as confirmed AD cases. We also stained for these 3 proteins in the locus coeruleus (LC), another brainstem region that has been implicated in early AD[16–18]. We then overexpressed a mutant form of human tau (P301L-tau) in the DRN of C57BL/6J mice and assessed behavior, 5-HT neuronal function, neuroinflammation, and the spread of tau pathology to interconnected brain regions. Together, these studies provide the first direct evidence that tau pathology in the DRN can induce depressive-like behavior, serotonergic dysfunction, and astrocytic activation in the absence of overt memory impairment. Furthermore, tau pathology that is initially confined to the DRN may spread to other brain regions, leading to the emergence of new symptoms.

## Results

### Tau pathology is present in the DRN of a subset of individuals without dementia

The presence of NFTs in the DRN has been reported in advanced AD [15, 19, 20], but only one study has reported this in individuals at an early stage[12]. We examined post-mortem brains from a group of 14 individuals without clinically diagnosed neurological disease (controls) and 10 AD cases. Tissues were sectioned through the brainstem and stained for tryptophan hydroxylase-2 (Tph2) as an anatomical reference for DRN and phospho-tau (ptau) (Ser202/Thr205) (AT8) to identify tau pathology. After eliminating samples without Tph2 immunoreactivity (Tph2-IR), 7 controls (65.43 ± 4.50 years; range: 54-80) and 10 AD cases (74.80 ± 3.86 years; range: 59-98) remained. We observed ptau-IR in the DRN of 57.14% (4/7) of control cases, which may indicate the DRN is one of the initial regions to develop tau pathology before an AD diagnosis. Among those individuals with diagnosed AD, 90% (9/10) had significant ptau-IR in the DRN (Table 1, Figure 1a-d, i).

**Figure 1:**
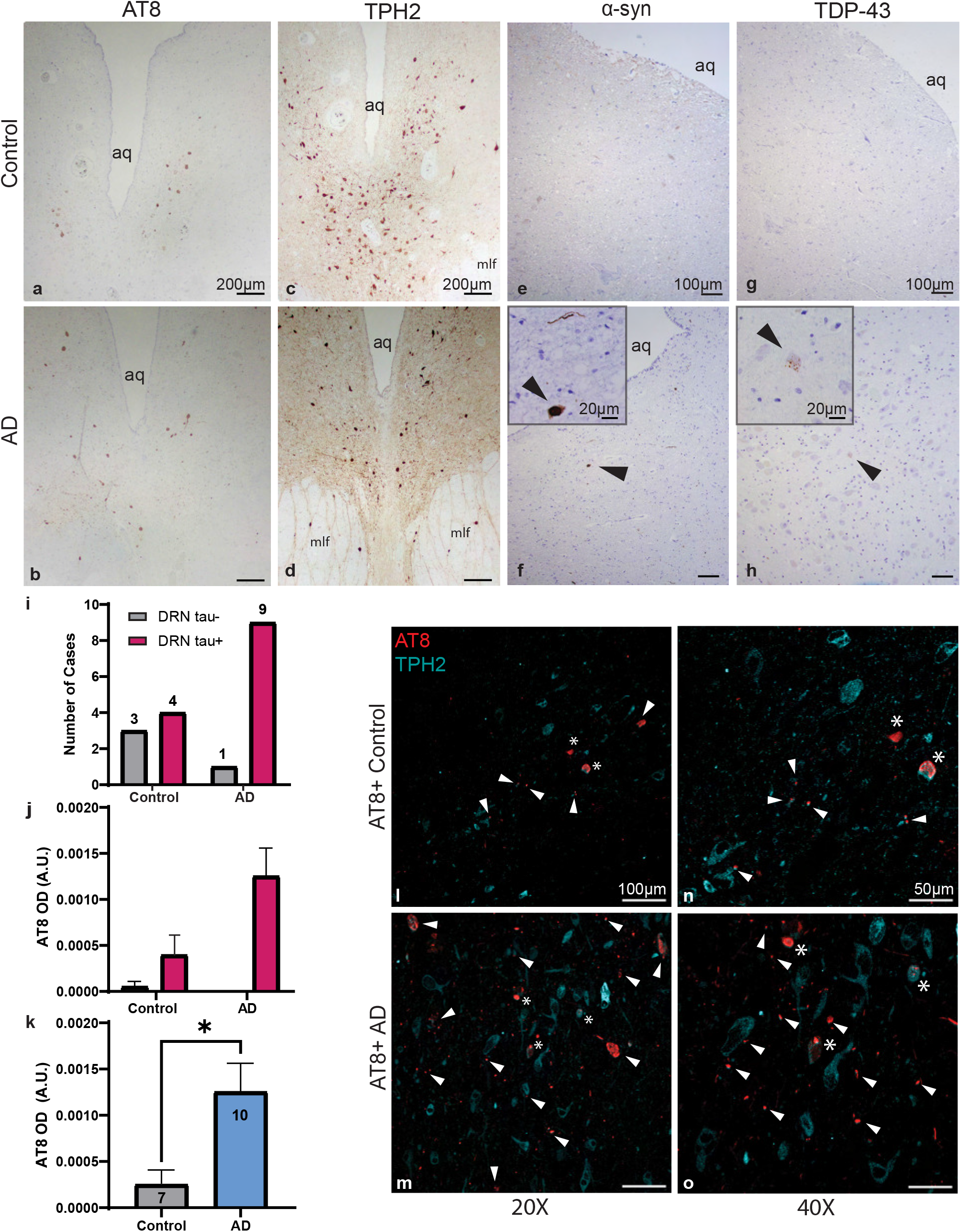
Neurodegenerative markers in the DRN of individuals cognitively normal and with Alzheimer’s Disease. Human dorsal raphe nucleus (DRN) sections were taken from post-mortem cases of individuals cognitively normal (Control) and diagnosed with AD at the time of death (AD). **a-b** DAB staining for AT8 (ptauSer202/Thr205) in control (a) and AD cases (b). **c-d** DAB staining of serotonergic cell marker Tryptophan hydroxylase 2 (TPH2) in control (c) and AD (d) cases, delineating the anatomical boundaries of the DRN. **e-f** DAB staining for α-syn (pS129-α-synuclein) in control (e) and AD (f) cases. Black arrowhead in f denotes α-syn+ staining, magnified in inset. **g-h** DAB staining for TDP-43 (pS409/410-TDP-43) in control (g) and AD (h). Black arrowhead in h denotes TDP-43+ staining, magnified in inset. Counterstain in a-h is hematoxylin. aq=cerebral aqueduct. mlf=medial longitudinal fasciculus. scale bars = 200 µm for a-d and 100 µm for e-h. For the insets of f-h, scale bar = 20um. **i** number of cases that are negative for (DRN tau-) and positive for (DRN tau+) tau in the DRN of control and AD groups. **j** Mean optical density values in control and AD cases, separated by presence or absence of tau staining. In a single tau-case among the AD group, OD=0 and does not appear on the axes. **k** Mean optical density values in control and AD cases, collapsed across DRN tau- and DRN tau+ cases. **l-m** Immunofluorescent staining of AT8 (red) and TPH2 (cyan) in tau+ control (l) and AD (m) cases taken at 20X magnification. **n-o** Magnified 40X images of l and m. White arrowheads point to colocalization of AT8 and TPH2 fluorescence. Asterisks denote larger regions of colocalization, such as cell bodies, that appear in magnified images l-o. Scale bar in l-m=100 µm, and 50 µm in n-o. Data are represented as mean ± SEM. \**p<*0.05.

**Table 1:** Post-mortem case demographics and dorsal raphe pathology.

| Case ID | Age | Sex | AD | Other ND | Ptau-DRN | a-syn DRN | TDP-43-DRN | Braak Stage | Source |
| --- | --- | --- | --- | --- | --- | --- | --- | --- | --- |
| 1 | 56 | M | No | No | - | - | - | NA | NIH |
| 2 | 54 | F | No | No | + | - | - | NA | NIH |
| 3 | 80 | F | No | No | + | + | - | III/IV | INC |
| 4 | 73 | F | No | No | - | - | - | V/VI | INC |
| 5 | 61 | M | No | No | + | - | - | 0 | INC |
| 6 | 80 | F | No | No | - | - | - | III/IV | INC |
| 7 | 54 | M | No | No | + | - | - | 0 | INC |
| 8 | 61 | F | Yes | No | + | - | + | V/VI | INC |
| 9 | 59 | M | Yes | No | - | - | - | V/VI | INC |
| 10 | 98 | F | Yes | No | + | - | - | V/VI | INC |
| 11 | 74 | M | Yes | LBD | + | + | - | V/VI | INC |
| 12 | 82 | M | Yes | LBD | + | + | - | III/VI | INC |
| 13 | 65 | M | Yes | No | + | - | + | V/VI | INC |
| 14 | 73 | M | Yes | No | + | - | - | V/VI | INC |
| 15 | 75 | M | Yes | LBD | + | + | - | V/VI | INC |
| 16 | 72 | M | Yes | No | + | - | + | V/VI | INC |
| 17 | 89 | F | Yes | No | + | - | - | V/VI | INC |

ND=Neurological disease
LBD = Lewy Body Disease
INC = Iowa Neurobank Core
NIH = National Institutes of Health
NA= Not Available

The DRN can also be affected in Lewy Body Dementia (LBD)[21], Parkinson’s disease (PD)[22], and frontotemporal lobar degeneration with TDP-43-immunoreactive pathology (FTLD-TDP)[23], so we also stained for pS129-α-synuclein and pS409/410-TDP-43 (Figure 1e-h). One control case was positive for α-synuclein in the DRN while none were TDP-43 positive. Three AD cases were positive for α-synuclein, all of which were comorbid for LBD. Another 3 AD cases, without α-synuclein pathology, showed positive staining for TDP-43. These data suggest that α-synuclein and TDP-43 pathology may be less prevalent in the DRN than tau. We then averaged the optical density of ptau staining across AT8- and AT8+ groups in the control and AD cases, (Figure 1j). To directly compare the control and AD groups, we took the average ptau OD for all control and AD cases and found a significant increase in the AD group (U=10, *p<*0.05), Figure 1k). Loss of serotonergic neurons is reported in advanced post-mortem AD cases [15, 24], so we next asked whether ptau colocalizes with Tph2 neurons in control and AD cases in our sample. Double immunofluorescence staining in DRN sections indicates that ptau does colocalize with a subset of Tph2-IR neurons in control and AD cases, but it is also found in other types of cells (Figure 1l-o).

Tau pathology in the LC has been described extensively and is noted to contain pre-tangle pathology prior to Braak Stage 1 [25]. Sections of the LC from the same cases were also assessed for dementia-related pathology, including ptau, pα-synuclein, and pTDP-43. LC samples were only available from 5 controls cases, 4 of which were ptau+. In the AD group, 9 out of 10 cases were ptau+. Of the 5 control cases that contained sections from both DRN and LC, 2 were positive for ptau in both regions, 2 had ptau in the LC only, and 1 had ptau in the DRN only. Of the 10 AD cases with both regions, 8 were positive for ptau in the DRN and the LC, 1 was positive for ptau in the DRN only, and 1 was positive for ptau in the LC only (Supplementary Figure 1a-b, e). It is worth noting that only one section was assessed per case for pathology, so it is possible that ptau pathology was missed in cases that contained tau only in the LC or DRN. One control case and 3 AD cases were also α-syn+ in both the LC and the DRN, all with comorbid LBD (Supplementary Figure 1c-d, f). Two out of these 3 AD cases that were positive for α-syn were also positive for ptau in the LC. We then asked whether tau or synuclein pathology in the LC colocalizes with noradrenergic neurons by performing a triple stain for tyrosine hydroxylase (TH), ptau, and p-αsyn. Here we observed colocalization between ptau and TH in a control and an AD case (Supplementary Figure 1g-h), as well as triple staining for ptau, α-syn and TH in an AD case (Supplementary Figure 1i). No cases were TDP-43+ in the LC (Table 2).

**Table 2:** Post-mortem case extended pathology and diagnoses.

| Case ID | Thal | CERAD | ADNC Score | DRN Threads | DRN Pre-tangles | DRN Tangles | ptau-LC | αsyn-LC | TDP43-LC | Diagnosis |
| --- | --- | --- | --- | --- | --- | --- | --- | --- | --- | --- |
| **1** | NA | NA | NA | 0 | 0 | 0 | NA | NA | NA | Control (ASCVD) |
| **2** | NA | NA | NA | 1 | 1 | 0 | NA | NA | NA | Control (ASCVD) |
| **3** | 2+ | 1 | Low/Int | 1 | 0 | 0 | + | + | - | Control (not specified) |
| **4** | 0 | 0 | Not | 1 | 0 | 0 | + | - | - | Control (COVID-19 pneumonia) |
| **5** | 0 | 0 | Not | 1 | 2 | 2 | + | - | - | Control (SARS-CoV-2 complications) |
| **6** | 2 | 0 | Int | 0 | 1 | 0 | + | - | - | Control (Acute bronchopneumonia) |
| **7** | 0 | 0 | Not | 1 | 1 | 0 | - | - | - | Control (Hypertensive cardiovascular disease) |
| **8** | 5 | 3 | High | 2 | 2 | 2 | + | - | - | AD (ADNC high) |
| **9** | 4 | 3 | High | 0 | 0 | 0 | + | - | - | AD (ADNC high) |
| **10** | 4 | 2 | High | 1 | 1 | 2 | + | - | - | AD (ADNC intermediate) |
| **11** | 4 | 2 | High | 2 | 2 | 1 | + | + | - | AD+LBD (ADNC high) |
| **12** | 3 | 2 | Int | 1 | 1 | 0 | - | + | - | AD+LBD (ADNC intermediate) |
| **13** | 5 | 3 | High | 1 | 1 | 2 | + | - | - | AD (ADNC high) |
| **14** | 5 | 2 | High | 2 | 2 | 1 | + | - | - | AD (ADNC intermediate) |
| **15** | 4 | 2 | High | 2 | 1 | 2 | + | + | - | AD+LBD (ADNC intermediate) |
| **16** | 5 | 2 | High | 1 | 1 | 1 | + | - | - | AD (ADNC high) |
| **17** | 5 | 2 | High | 1 | 1 | 0 | + | - | - | AD (ADNC intermediate) |

NA= Not Available
Int = Intermediate
ASCVD = Atherosclerotic cardiovascular disease
ADNC = Alzheimer's disease neuropathological change
0: no pathology ;1: rare pathology; 2: frequent pathology. Control cases 4 and 6 exhibited extremely rare pathology, and were assigned a score above 0.

### Overexpression of P301L-tau in the DRN promotes depressive-like behaviors in mice

We then asked whether tau pathology in the DRN could induce behavioral impairments associated with prodromal AD. An AAV vector expressing the human P301L-tau mutant or GFP (control) was stereotaxically injected into the DRN of male C57BL/6J mice at 8 weeks of age. Behavioral testing began 4 weeks later. C57BL/6J mice express endogenous mouse tau which does not form tau aggregates in the absence of human tau, so significant tau pathology was only expected in the mice injected with P301L-tau in the DRN (Figure 2a-c). We found that P301L-tau^DRN^ mice exhibited a robust deficit in social interaction relative to controls (F_1,18_=64.97, *p<*0.0001, stranger x tau interaction). Bonferroni post-tests revealed that the presence of DRN P301L-tau significantly reduced time spent with the stranger (t_36_=6.49, *p<*0.0001) and increased time spent with the empty cage (t_36_=4.14, *p<*0.001) (Figure 2d). There was also a reduction in sucrose preference on the test day (t_17_=2.92, *p<*0.01) (Figure 2e-f). The two groups did not differ significantly in immobility time in the forced swim test (t_18_=1.056, ns), which is a putative measure of depressive-like behavior (Figure 2g). However, because the test relies on immobility as the dependent variable, heightened locomotor activity in P301L-tau^DRN^ mice may have obscured depressive-like behaviors in this test.

**Figure 2:**
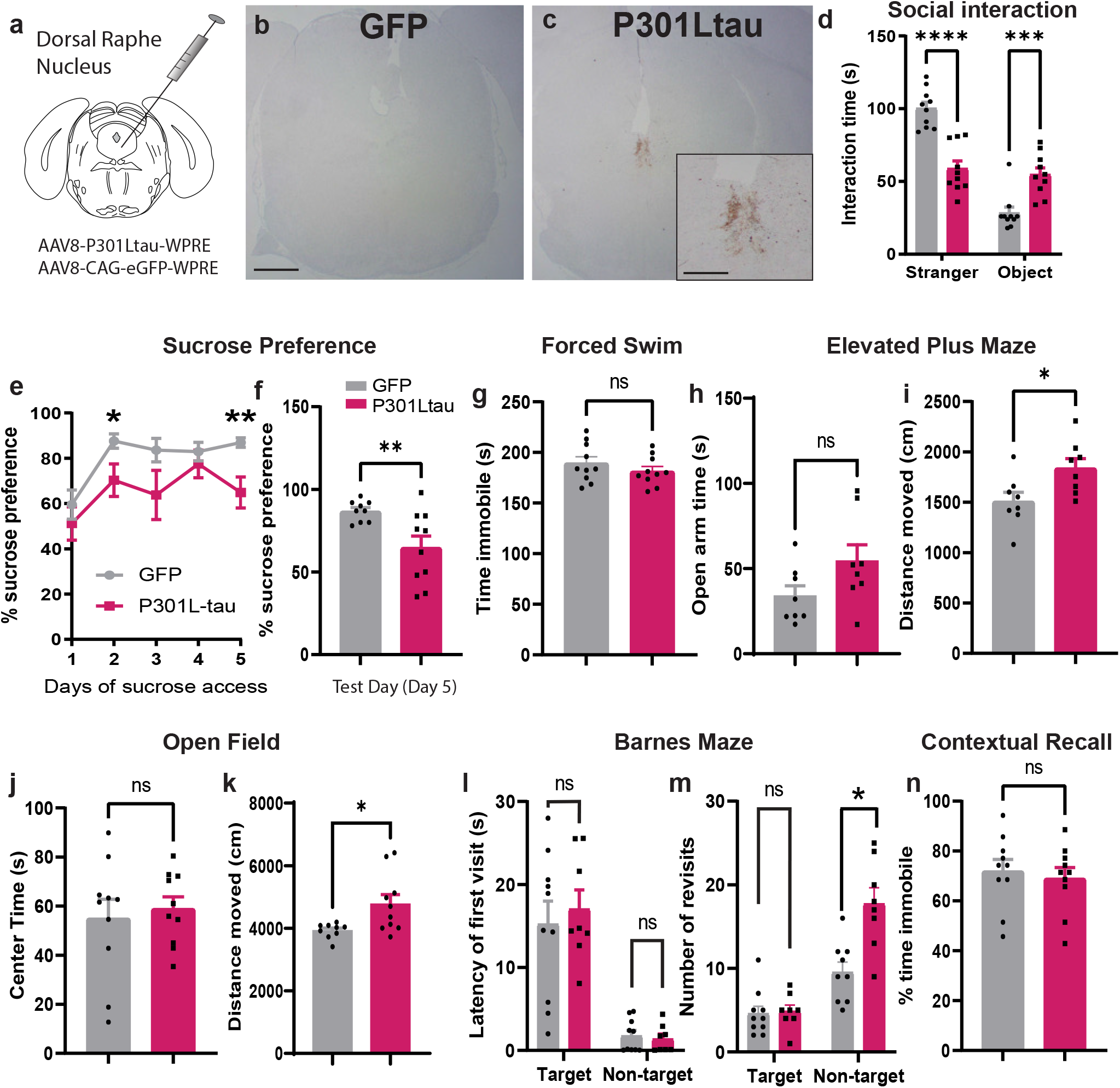
DRN tau pathology promotes depressive-like behaviors in male mice. The P301L-tau mutation or green fluorescent protein (GFP) was delivered via adeno-associated virus into the DRN of C57BL/6J mice (n=10 per group, 20 total). **a** Schematic of viral infusion site in the DRN. **b-c** Representative DAB images of viral P301L-tau placement staining in GFP (b) and P301L-tau animals (c). **d** Graph of mean time GFP (gray) and P301L-tau mice (magenta) spent interacting with a stranger mouse and object in the social interaction assay. **e** X-Y plot of percent sucrose preference across all days of the sucrose preference assay. **f** Bar graph of percent sucrose preference on test day for both groups (day 5). **g** Bar graph depicting the percent of time mice spent immobile in the forced swim assay. **h-i** Time spent in the open arms (h) and distance moved (i) in the elevated plus maze (EPM). **j-k** Time mice spent in the center of the arena (j) and distance moved (k) in the open field test (OFT). **l-m** Latency of first visit (l) and number of revisits (m) to target and non-target holes in the Barnes Maze task. **n** Percent time mice spent immobile in contextual fear conditioning task. Data are represented as mean ± SEM. Scale bar= 500 µm in (b) and 200 µm in (c). \**p<*0.05, \*\**p<*0.01, \*\*\**p<*0.001, \*\*\*\**p<*0.0001.

The elevated plus maze (EPM) behavioral test is commonly used to assess generalized anxiety-like behaviors in rodents. Here we did not observe significant group differences in time spent in the open arms (Figure 2h), although there was a trend toward an increase (t_14_=1.85, *p=*0.085). There was, however, a significant increase in locomotor activity in the EPM (t_14_=2.56, *p<*0.05) (Figure 2i). In the open field test, we also observed an increase in locomotor activity in the first 10 minutes (t_10.28_=2.75, *p<*0.05, Welch’s correction), but there was no change in time spent in the center of the arena (t_17_=0.51, ns) (Figure 2j-k). These results indicate that P301L-tau in the DRN may increase locomotion without altering anxiety-like behavior. We next examined spatial memory in the Barnes Maze and found no evidence of deficits in latency to the target hole (t_16_=0.50, ns) (Figure 2l) or the frequency of target revisits (t_16_=0.24, ns) (Figure 2m). There was an increase in the frequency of non-target visits (t_16_=2.37, *p<*0.05), although the latency to the first non-target hole (t_16_=0.43, ns) did not change. Similarly, there was no spatial memory deficit in the contextual fear recall test (t_18_=0.46, ns) (Figure 2n). These results suggest that while these mice display a depressive-like behavioral phenotype as well as hyperlocomotion, spatial memory is still intact.

Sex differences have been reported in the severity of AD neuropathology, with women typically having more NFTs, greater cognitive decline, and accelerated disease progression relative to men [26–29]. The DRN is a sexually dimorphic region that is enriched in both types of estrogen receptors (ERα and Erβ)[30], so we expected to observe sex differences in behavioral manifestations of DRN neuropathology. Here we observed a significant reduction in social interaction in female P301L-tau^DRN^ mice relative to controls (F_1,28_=24.79, *p<*0.0001, stranger x tau interaction). Bonferroni post-tests also indicated a significant reduction in time spent with the stranger (t_56_=5.11, *p<*0.0001), but not with the empty cage (t_56_=2,15, ns) (Supplementary Figure 2a-b). However, unlike the males, there was no change in sucrose preference in females (t_28_=0.464, ns) (Supplementary Figure 2c-d). There were also no differences in anxiety-like behavior in the EPM (t_28_=0.424, ns) or open field (t_28_=1.06, ns), nor any change in locomotor activity in either test (EPM: t_28_=0.450, ns; OF:t_28_=1.15, ns) (Supplementary Figure 2e-h). We then tested for spatial memory deficits in the contextual fear recall test and found no differences between the two groups (t_28_=0.104, ns) (Supplementary Figure 2i). Contrary to our expectations, these results indicate that female P301L-tau^DRN^ mice have fewer behavioral impairments relative to controls than males. However, the accelerated rate of decline in women with AD may be related to a post-menopausal reduction in estrogen and progesterone[31, 32], so these differences may only begin to emerge in aged female mice.

### Tau pathology in P301Ltau^DRN^ mice is predominantly found in neurons

The introduction of P301L-tau in the DRN is expected to increase ptau staining in regions of viral dispersion (Figure 3a-d). We first quantified ptau-Ser202/Thr205 (AT8) optical density (OD) across subregions of the DRN (rostral, mid, caudal) and found a significant main effect of P301L-tau injection (F_1,36_=16.61, *p<*0.001), but no interaction between tau and subregion. Bonferroni post-tests revealed a significant increase in the rostral DRN (t_8.4_=3.54, *p<*0.05) in P301L-tau^DRN^ mice relative to GFP^DRN^ mice in an anatomically matched area. This increase in ptau persisted when slices were pooled across subregions (t_8.37_=3.03, *p<*0.05; Welch’s correction) (Figure 3e). The DRN contains 5-HT neurons that are vulnerable to neurodegeneration in AD and may be sensitive to the presence of tau tangles. We next examined Tph2 neuronal density and OD in regions of interest (ROIs) that contained AT8 staining. Surprisingly, there was no change in the density of Tph2-expressing neurons in any subregion of the DRN of P301L-tau^DRN^ mice (F_1,34_=0.168, ns) or pooled across subregions (t_11.37_=0.060, ns, Welch’s correction) (Figure 3f). Tph2 OD was also unchanged in all subregions (F_1,34_=0.815, ns) and when pooled across subregions (t_16_=1.06, ns) (Figure 3g). However, we did observe a negative correlation between Tph2 cell density and AT8 OD in AT8+ regions of the DRN in P301L-tau mice (R^2^=0.4707, *p<*0.05) (Figure 3h). The direction of this correlation suggests that there may be a reduction in Tph2 as tau pathology accumulates in the P301L-tau group. Although P301L-tau expression was not restricted to 5-HT neurons in this study, a notable percent of Tph2 neurons were also AT8+ within regions of AT8 immunoreactivity (Rostral: 34.14 ± 5.56 %, Mid: 35.98 ± 8.23 %, Caudal: 26.21 ± 7.06 %) (Figure 3i).

**Figure 3:**
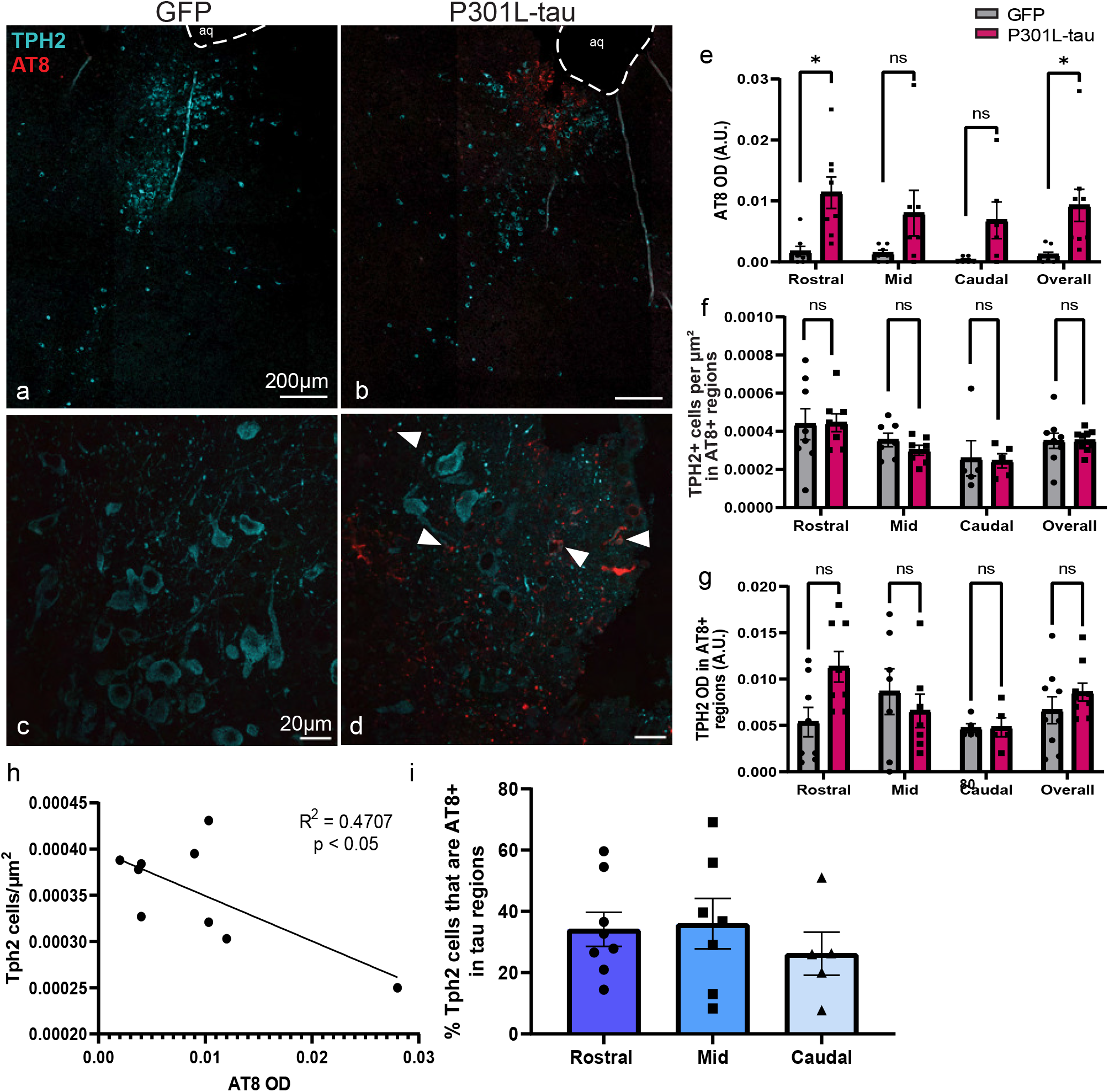
Tau pathology colocalizes with serotonergic DRN neurons. **a,c** Representative images of TPH2 (cyan) and AT8 (red) immunofluorescent staining in the DRN of a mouse that received AAV-GFP at 15X and 40X, respectively. **b,d** Representative images of a mouse that received AAV-P301L-tau, also at 15X and 40X. White arrowheads point to instances of TPH2 and AT8 colocalization. **e** Bar graph of AT8 optical density (OD) of GFP (gray) and P301L-tau (magenta) animals across subregions of the DRN and pooled across subregions (overall). **f** Bar graph of TPH2+ cell density in ROIs that contained AT8+ staining. **g** Graph of TPH2 OD in ROIs that were AT8+. We observed no differences between GFP and P301L-tau mice. **h** Simple linear regression between TPH2+ cell density and AT8 OD in P301L-tau animals. **i** Bar graph of the percentage of TPH2+ cells that were also AT8+ in ROIs for P301L-tau animals.

Since P301L-tau delivery was not restricted to a particular population of cells, we next determined the relative abundance of tau pathology in neurons and glia. Again, using regions of AT8 immunoreactivity, we calculated the % of AT8 immunoreactive objects (including cell bodies and puncta in dendrites and axons) co-positive for one of 3 general cell-type markers: NeuN for neurons, GFAP for astrocytes, and Iba1 for microglia at 4 weeks and 8 weeks post-viral injection (Figure 4). At both 4 and 8 weeks after viral injection, we observed that AT8 was most frequently co-localized with NeuN (27.81 ± 5.17 % of AT8+ staining was co-labeled for NeuN at 4 weeks, 40.06 ± 2.364 % at 8 weeks), with a notable increase in ptau-NeuN colocalization over time. We also noted what appears to be a shift in the proportion of AT8 that co-localized with Iba1 from 4 to 8 weeks of viral expression: 11.01 ± 3.96 % was Iba1+ at 4 weeks, decreasing to 9.71 ± 1.51 % at 8 weeks; as well as a slight increase in AT8-GFAP+ areas from 4 to 8 weeks: 9.28 ± 2.58 % at 4 weeks and 11.60 ± 2.31 at 8 weeks (Figure 4e-f, k-l).

**Figure 4:**
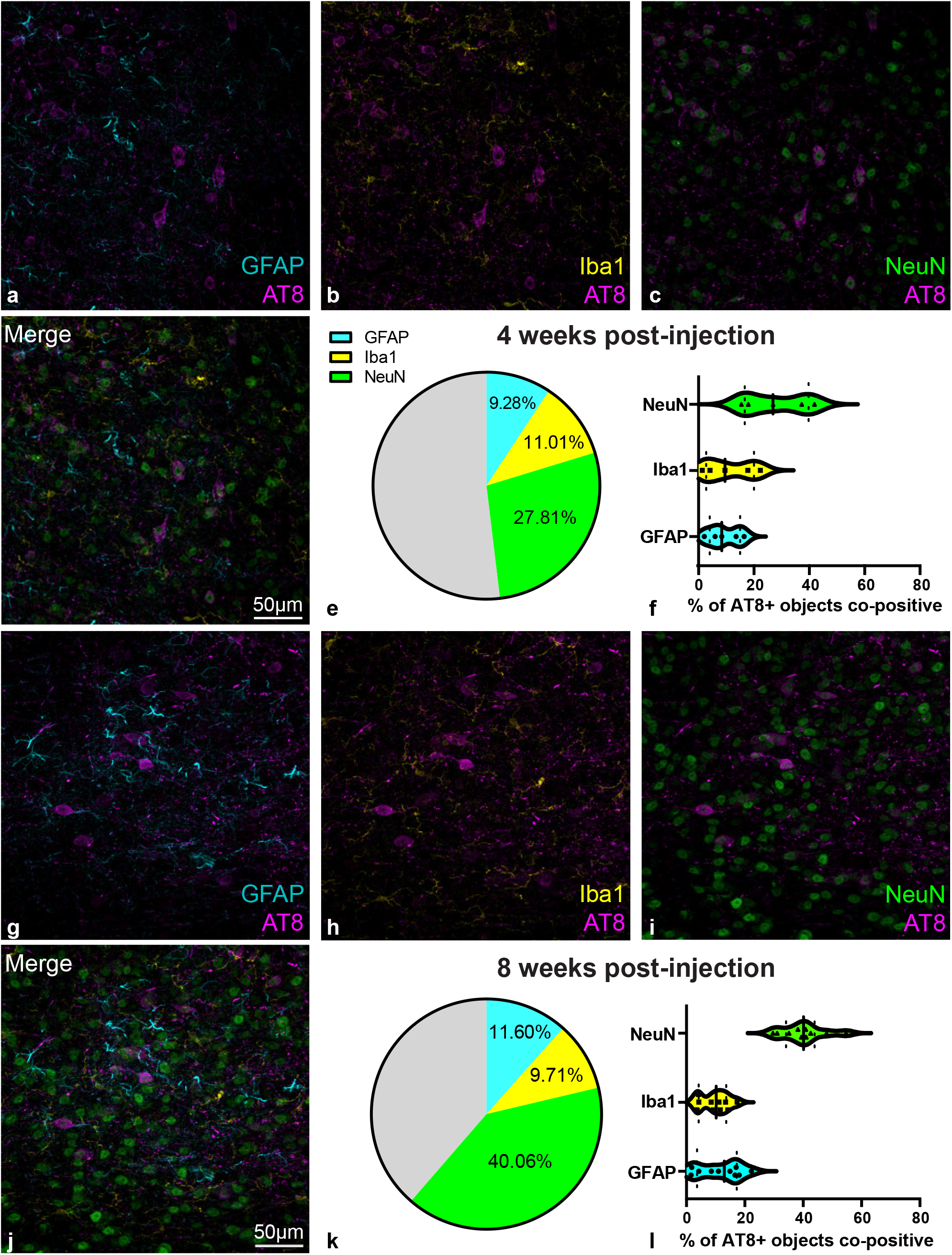
Tau pathology preferentially colocalizes with neurons instead of glia in the DRN. Representative images from C57BL/6J mice injected with AAV-P301L-tau into the DRN after 4 weeks (a-d) and 8 weeks (g-j) of viral transduction. **a,g** Images showing AT8 (magenta) and GFAP (cyan) immunflourescence staining. **b,h** Images showing AT8 and Iba1 (yellow) staining. **c,i** Images of AT8 and NeuN (green). **d,j** Combined views of the AT8-GFAP-Iba1-NeuN IF depicted in a-c and g-i. White arrows point to areas of colocalization. Images taken at 40X. Scale bar = 50 µm. **e,k** Pie chart depicting the percent of AT8+ staining that was colocalized with GFAP (cyan), Iba1 (yellow), or NeuN (green). Gray indicates AT8-only, presumably in processes not positive for one of the 3 cell markers. **f,l** Violin plots showing median % positive staining (black line), quartiles (dashed lines), and distribution of data points used in the % calculations.

### Tau pathology activates local astrocytes and induces inflammatory gene expression in the DRN

We then investigated whether P301L-tau expression in the DRN altered astrocytic and microglial markers in AT8 immunoreactive areas of the DRN (Supplementary Figure 3a-d). Here we found a significant increase in GFAP+ OD (t_15_=2.65, *p<*0.05) and GFAP+ astrocytic cell density in P301L-tau^DRN^ vs GFP^DRN^ controls (t_11.31_=2.88, *p<*0.05; Welch’s correction) (Supplementary Figure 3e-f). However, Iba-1 optical density (t_16_=1.11, ns) and Iba-1+ microglial cell density (t_16_=0.344, ns) did not change. This increase in astrocytic activity may be indicative of neuroinflammation which could account for the hyperlocomotion exhibited by P301L-tau^DRN^ mice. This is congruent with a previous study in which IL-1β overexpression in the DRN induced hyperlocomotion or “mania-like behavior” in the EPM [33]. Surprisingly, when we examined gene expression of neuroinflammatory-related genes, there was no change in IL-1β mRNA. There was in increase in IL-1α which activates a subtype of reactive astrocytes and induces them to generate reactive oxygen species (ROS) that promote degeneration[34] (t_10.67_=3.58, *p<*0.01; Supplementary Figure 3g). Interestingly, IL-12β, a pro-inflammatory cytokine that mediates host-pathogen interactions and has anti-tumor activity activity[35], was reduced in P301L-tau^DRN^ mice (t_7.05_=2.69, *p<*0.05). In a recent study, a single-nucleotide polymorphism (SNP) in IL-12β linked to various types of cancer was also found to be associated with a heightened risk of late-onset AD [36]. Finally, we observed an increase in Fyn-related kinase (Frk) (t_8.82_=2.28, *p<*0.05) which was elevated in the DRN of mice expressing wild-type human tau[37] and is known to contribute to the phosphorylation of tau.

### Tau pathology increases 5-HT neuronal excitability and glutamatergic transmission

We next asked whether the presence of tau pathology in the DRN altered the intrinsic excitability of 5-HT neurons in the DRN (Figure 5a-b). The action potential threshold, or rheobase, was significantly lower in P301L-tau^DRN^ mice (t_27_=2.17, *p<*0.05), which is suggestive of hyperexcitability (Figure 5c-d). There was also a shift towards a more depolarized resting membrane potential (RMP) (-57.81 ± 1.71 mV for GFP and -52.38 ± 1.67 mV for P301L-tau, t_27_=2.24, *p<*0.05) as well as and an increase in the input resistance (t_27_=2.36, *p<*0.05) which may account for the reduced rheobase (Figure 5e-f). We then looked at action potential frequency at 10 pA current steps from 0-200 pA. This revealed a significant increase in current-induced spiking (Tau x current interaction: F_20,540_=2.71, *p<*0.0001; main effect of tau: F_1,27_=4.82, *p<*0.05). Bonferroni *post hoc* tests showed a significant increase at 180 pA (t_567_=3.08, *p<*0.05), 190 pA (t_567_=3.08, *p<*0.05), and 200 pA steps (t_567_=3.24, *p<*0.05) (Figure 5g). Next, we repeated the current-step protocol from a starting potential of -70 mV to adjust for variations in RMP. Here we also observed a significant increase in action potential frequency as a function of current (Tau x current interaction: F_20,520_=3.75, p>0.0001; main effect of tau: F_1,26_=4.90, *p<*0.05), and Bonferroni *post hoc* tests also indicated an increase at the 180 pA (t_546_=3.21, *p<*0.05), 190 pA (t_546_=3.51, *p<*0.05) and 200 pA steps (t_546_=3.22, *p<*0.05). (Figure 5h).

**Figure 5:**
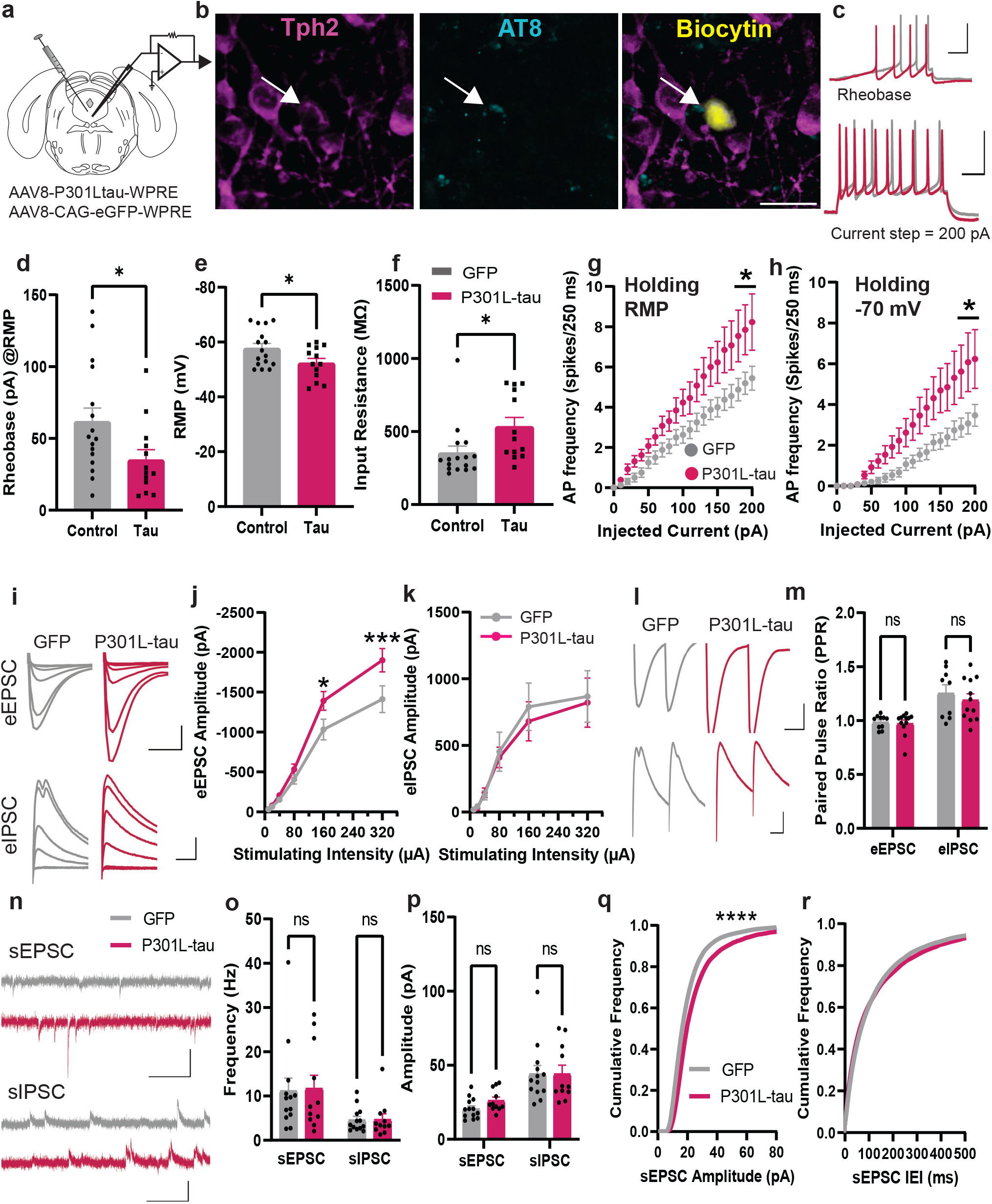
Neuronal excitability and glutamatergic transmission is elevated in P301L-tau^DRN^ mice. **a** Stereotaxic injection of AAV encoding P301L-tau and GFP into the DRN. **b** Representative confocal image of recorded neuron containing biocytin. *Left:* Tph2 immunoreactivity (magenta) with recorded neuron indicated with a white arrow. *Center:* AT8 immunoreactivity (cyan) with recorded neuron indicated with a white arrow. *Right:* Merged image containing Tph2, AT8, and biocytin (yellow). **c** *Top:* Representative traces of action potential firing during a current ramp protocol used to compute rheobase (x-axis = 200 ms, y-axis = 50 mV). *Bottom:* Representative traces showing action potential firing during a 200 pA current step (x-axis = 50 ms, y-axis = 50 mV). **d-f** Histogram of mean action potential thresholds (rheobase), resting membrane potential (RMP), and input resistance for control and P301L-tau^DRN^ mice. **g-h** Histogram of action potential frequency as a function of current (0-200 pA) starting at RMP and at -70 mV. **i** *Top:* Representative traces of eEPSCs at a range of stimulation intensities (10 - 320 µA) (x-axis = 10 ms, y-axis = 500 pA). *Bottom:* Representative traces of eIPSCs (x-axis = 10 ms, y-axis = 200 pA). **j-k** Histogram of mean eEPSC amplitude and amplitude as a function of stimulation intensity. **l** Representative traces of PPR for eEPSCs (x-axis = 10 ms, y-axis = 500 pA) and eIPSCs (x-axis = 10 ms, y-axis = 200 pA). **m** Histogram of mean PPR for eEPSCs and eIPSCs. **n** Representative traces of sEPSCs (x-axis = 100 ms, y-axis = 50 pA) and sIPSCs (x-axis = 200 ms, y-axis = 50 pA) in control and P301L-tau^DRN^ mice. **o-p** Histograms of mean frequency and amplitude of sEPSCs and sIPSCs. **q-r** Cumulative frequency distributions of sEPSC and inter-event intervals (IEIs) in GFP and P301L-tau^DRN^ mice. Data are represented as mean ± SEM unless otherwise indicated. Scalebars = 100 µm. \**p<*0.05, \*\*\**p<*0.001, \*\*\*\**p<*0.0001.

The DRN contains glutamatergic neurons and receives glutamatergic inputs from outside the DRN that may take up tau from the extracellular space. Both could affect excitatory drive in 5-HT neurons. Alternatively, intracellular signaling cascades on the postsynaptic side may be impacted by the presence of tau pathology in 5-HT neurons. To address this question, we recorded evoked excitatory post-synaptic currents (eEPSCs) in 5-HT neurons and found an increase in eEPSC amplitude (Tau x stimulation intensity interaction: F_5,100_=4.46, *p<*0.01; main effect of tau: F_1,20_=4.52, *p<*0.05). *Post hoc* Bonferroni comparisons were significant at the 160 pA (t_120_=2.99, *p<*0.05) and 320 pA stimulation intensities (t_120_=4.08, *p<*0.001) (Figure 5i-j). There is also a population of GABAergic interneurons that provide inhibitory input onto 5-HT neurons in the DRN, so any change in the function of these neurons could alter synaptic drive as well. However, we did not see group differences in evoked inhibitory postsynaptic current (eIPSC) amplitude (Tau x stimulation intensity interaction: F_5,95_=0.1334, ns; main effect of tau F_1,19_=0.080, ns) (Figure 5k). We then calculated the paired pulse ratio (PPR) of eEPSCs to determine whether the presence of tau could alter the probability of transmitter release at glutamatergic synapses onto DRN 5-HT neurons, but we did not see any group differences (t_20_=0.3532, ns). Likewise, there was no change in PPR of eIPSCs (t_19_=0.6804, ns), suggesting no change in the probability of GABA release (Figure 5l-m).

We next turned our attention to spontaneous EPSC and IPSCs (sEPSCs and sIPSCs respectively), which are the result of action potential-independent neurotransmitter release. Neither sEPSC nor sIPSC frequencies were altered (EPSC freq: t_22_=0.1294, ns; IPSC freq: t_22_=0.0832, ns), suggesting no change in presynaptic transmitter release (Figure 5n-o). We did see a trend toward an increase in sEPSC amplitude, which is consistent with our eEPSC results (t_22_=1.90, *p=*0.07), but no change in sIPSC amplitude (t_22_=0.0011, ns) (Figure 5p). We did see a significant rightward shift in the cumulative frequency distribution curve of sEPSC amplitude (Kolmogorov-Smirnov D=0.1454, *p<*0.0001), suggesting that P301L-tau mice tend to have a higher number of sEPSCs with larger amplitudes (Figure 5q). On the other hand, the cumulative frequency distribution of the inter-event interval (IEI) of sEPSCs did not differ between groups (K-S D=0.0708, ns), once again indicating no change in sEPSC frequency in P301L-tau^DRN^ mice (Figure 5r).

### Propagation of tau pathology from the DRN to other brain regions

Tau pathology propagates from one neuron to another by a process known as transsynaptic spread. When initially confined to the entorhinal cortex of C57BL/6J mice by stereotaxic injection of a viral vector encoding P301L-tau, tau pathology was reported to spread to the hippocampus and impair hippocampal-dependent LTP[38]. Likewise, tau fibrils injected into the LC of PS19 mice were found to propagate tau pathology to LC projection areas, as well as sources of LC innervation in the brain, including the hypothalamus, amygdala, bed nucleus of the stria terminalis, and frontal cortex [39]. We examined AT8 staining in areas that are known targets of the DRN 5-HT system, as well as regions that develop tau pathology in the early stages of AD in a subset of P301L-tau^DRN^ mice at approximately 2 months post-injection via DAB staining. These areas included the entorhinal cortex (EC), thalamus (Thal), hippocampus (HP), hypothalamus (HT), amygdala (AMY), and caudate/putamen (CPu) (Figure 6a-f). Our results suggest that tau pathology can spread from DRN to the thalamus, hypothalamus, and amygdala. One of the mice also had AT8 staining in the CPu, while none had evidence of tau pathology in the entorhinal cortex or hippocampus (Figure 6g). It should be noted that the tau pathology was mostly confined to axon terminals and was not detected in neuronal perikarya. We then asked whether tau pathology that had migrated to downstream regions colocalized with serotonergic axons using immunofluorescence (IF). In a subset of brains 4 weeks and 8 weeks post-injection of P301L-tau in the DRN, we observed human tau staining (HT7) in the same regions identified by the DAB stain and determined that these projections colocalized to varying degrees with SERT (Suppl. Figure 4). Across all regions sampled for analysis, 24.47 ± 5.13 % of the tau+ IR was also positive for SERT 4 weeks after P301L-tau injection (Suppl Fig 4c) and decreased to 11.72 ± 3.69 % by 8 weeks. The general trend of decreased in tau-SERT colocalization held true across subregions, except in the amygdala and hippocampus (Suppl Fig 4c-d). The sample size here was powered to verify the presence or absence of any colocalization, which we did confirm. Further research with a larger sample would be needed to verify these estimates of the degree of colocalization. Suppl Table 3 has a comprehensive list of regions where tau was observed via IF.

**Figure 6:**
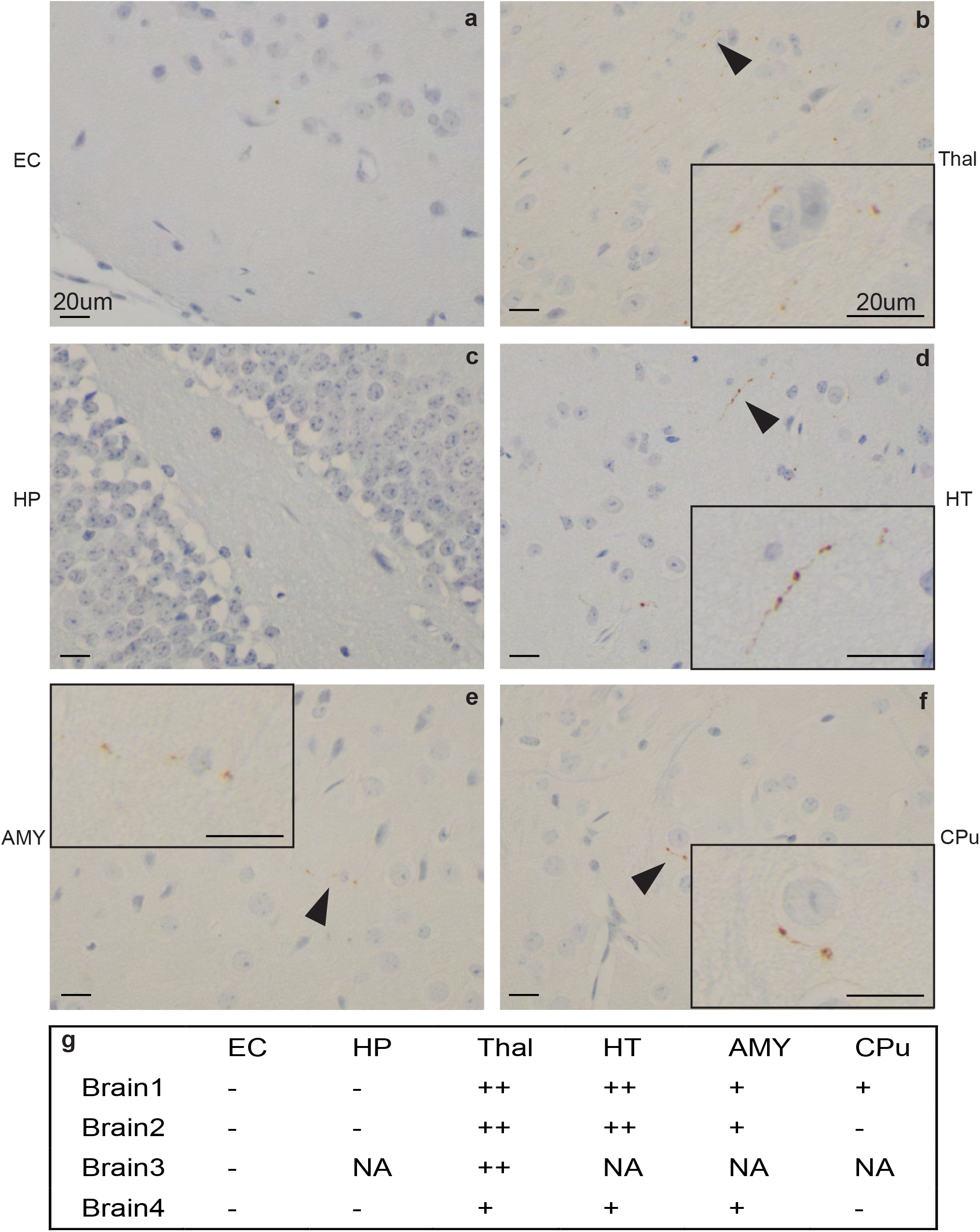
Tau pathology propagates from the DRN to other brain regions. In a subset of male mice (n=4), we evaluated brain regions downstream of the DRN known to be involved in affective changes and memory for AT8 staining. **a-f** Representative images of staining in respective regions at 60X objective magnification. Insets in **b, d-f** are enlarged and cropped from larger images to highlight positive staining. Black arrowheads point to AT8+ staining. **g** Table summarizing semi-quantitation of AT8 staining in downstream regions. NA=no sections available of the region from that brain, (-) indicates no AT8 staining. (+) indicates very little pathology localized to single fields of view within a region. (++) indicates multiple areas in multiple fields of view with AT8 staining in a region. Scale bars=20 µm. EC=entorhinal cortex, Thal = thalamus, HP= hippocampus, HT=hypothalamus, AMY=amygdala, CPu=caudate/putamen

## Discussion

Alzheimer’s disease is a fatally progressive neurodegenerative disease that will affect an estimated 8.4 million Americans by 2030[28]. It is often diagnosed at an advanced stage where significant neurodegeneration and cognitive impairment has occurred, rendering most treatments inefficacious. Early detection and intervention are imperative to halt the progression of neuropathology before widespread neurodegeneration occurs, but it requires predictive biomarkers and diagnostic criteria that can be implemented in the clinic. Establishing early biomarkers of AD that proceed neurodegeneration by several years or decades would represent a breakthrough for AD diagnosis and prevention. Our data strongly suggest that tau pathology in the DRN may cause behavioral or mood disruptions that manifest early in AD, well in advance of any overt cognitive symptoms. This is supported by several other studies in the literature indicating that subcortical nuclei are affected by tau pathology prior to involvement of the entorhinal cortex[12, 40–42]. Although we cannot know for certain whether these individuals without cognitive impairment would have gone on to develop dementia, our behavioral data in P301L-tau^DRN^ mice suggests that the presence of tau pathology in the DRN can promote depressive-like behavior which is one of the hallmarks of prodromal AD. These behavioral changes coupled with serotonergic dysfunction, astrocytic activation, and tau propagation to other brain regions are all consistent with an early AD profile.

One of the main limitations of this study is that appropriate sections of the DRN are often not available from banked brains, so our post-mortem human data are based on a relatively small number of cases. Despite standardized protocols for brain banking, the small size and cytologic diversity of the DRN means that the level of section is difficult to standardize, precluding accurate assessment of 5-HT neuronal density. However, we were able to assess Tph2 expression and cell density in the mice injected with P301L-tau in the DRN. The finding that Tph2-expressing neurons remained relatively intact in P301L-tau^DRN^ mice was unexpected, as 5-HT neuronal depletion in AD post-mortem tissue has been reported elsewhere[15]. There may be some compensatory upregulation of Tph2 in cells with mild to moderate tau accumulation, with Tph2 depletion only occurring in the later stages as tau pathology progresses. This is supported by the fact that there is a negative correlation between Tph2 cell density and AT8 OD in the DRN of P301L-tau^DRN^ mice. We also note that P301L-tau is associated with FTD/Parkinsonism, and future directions will include other types of tau pathology to determine if there are similar effects in the DRN.

We did observe an increase in the intrinsic excitability and enhanced glutamatergic transmission in P301L-tau^DRN^ mice. The increased evoked EPSC amplitude in the absence of any change in PPR suggests a post-synaptic mechanism, which may be due to altered glutamate receptor expression, trafficking, or intracellular signaling. Excessive activity in 5-HT neurons may have accounted for some of the behavioral dysregulation we observed here, including hyperactivity and depressive-like behaviors. In a previous study by Ren and colleagues, different subgroups of 5-HT neurons were shown to activate distinct and opposing behavioral programs depending on their downstream projection patterns, and 5-HT neurons that project to subcortical regions tended to promote aversive behaviors[11, 43]. While we did not target specific 5-HT subpopulations in this study, it is possible that hyperactivity in subcortical-projecting 5-HT neurons could underpin the behavioral phenotypes observed here. There is also evidence to suggest that heightened DRN serotonergic activity could underlie some depressive disorders, with depressed suicidal patients exhibiting elevated Tph2 expression in the DRN and its projection areas[44–49].

Hyperactivity in 5-HT neurons with tau pathology may seem paradoxical, but there is evidence to suggest that it may be an early indicator of neuronal dysfunction in AD and a precursor to neurodegeneration[50, 51]. The higher protein load puts stress on the cell, so this hyperactivity may be compensatory to maintain network performance in the face of increased energy demands. Once the pathology progresses beyond a certain critical point where the cell can no longer compensate, there may be a subsequent reduction in activity. Another intriguing possibility relates to the marked increase in astrocytic density and GFAP immunoreactivity in P301L-tau^DRN^ mice, which is a well-known hallmark of AD[52]. Astrocytes usually proliferate in response to brain injury or neuroinflammation and can assume a neurotoxic A1 or neuroprotective A2 phenotype[53]. When activated, astrocytes also modulate neuronal excitability by changing the concentration of K+ ions in the extracellular space, a process known as K+ clearance[54]. Any disruption in these astrocytic K+ clearance mechanisms can result in aberrant neuronal activity, such as the hyperexcitability we observed here. Astrocytes also play a central role in glutamate homeostasis in the CNS, so any perturbations in glutamate uptake or release by local astrocytes could alter the excitatory/inhibitory balance in surrounding neurons and lead to glutamate excitotoxicity[55]. We did observe an increase in sEPSC amplitude in 5-HT neurons, but not frequency, suggesting that synaptic glutamate release is not affected but there may be dysregulation of astrocytic glutamate release or uptake. In cultured hippocampal neurons, it was reported that tau accumulation in astrocytes disrupted Ca^2+^-dependent gliotransmitter release, which in turn altered glutamate transmission in surrounding neurons[56]. P301L-tau is known to accumulate in both neurons and glia[57], and frontotemporal lobar degeneration (FTLD-tau/P301L) is characterized by the presence of tau-immunoreactive deposits in astrocytic processes [57, 58]. Although P301L-tau accumulation in the DRN was primarily neuronal, ∼12% of tau pathology in the DRN colocalizes with astrocytes, which may have altered their K+ buffering and glutamate uptake capacities, causing a shift toward hyperexcitability in 5-HT neurons.

Astrocytes are also activated in the presence of pro-inflammatory cytokines released by microglia in response to tau pathology, resulting in the neurotoxic reactive A1 phenotype [59–62]. Although we did not observe a change in microglial density in the DRN, we did see an increase in the pro-inflammatory cytokine IL-1α which activates A1 astrocytes[34]. Alternatively, astrocytes can also be neuroprotective. A study in human subjects with tau pathology indicates that those without dementia contained astrocytes with upregulated glutamate transporter 1 (GLT-1) expression compared to those with dementia, suggesting that GLT-1 may play a neuroprotective role in these astrocytes[63]. In a mouse model of human tau that was under the control of a GFAP promoter and exclusive to astrocytes, downregulation of GLT-1 preceded tau accumulation[64]. It would be an interesting future direction to quantify GLT-1 expression within astrocytes in P301L-tau^DRN^ mice alongside other neuroprotective markers.

Tau propagation via transneuronal spread is well-documented in the literature and can be accelerated by microglial activation and neural activity[65–67]. There is also emerging evidence that astrocytes can participate in the spread of tau to other astrocytes or even accelerate transneuronal spread[68, 69]. Our results confirm that tau pathology initially confined to the DRN can propagate to the thalamus, hypothalamus, and amygdala and appears to colocalize with some serotonergic processes. While tau spread was restricted to axonal processes, we think it is possible that tau pathology may accumulate in downstream targets over time and eventually be taken up by adjacent cell bodies. The amygdala and hypothalamus have both been implicated in depressive behaviors, but the accumulation of tau in these areas was sparse and not likely to be a significant driver of the behavioral phenotypes observed here in P301L-tau^DRN^ mice. It is also worth noting that tau spread to the entorhinal cortex and hippocampus was very sparse even after 8 weeks, so the DRN may not be a major source of tau pathology in those areas in the early stages of AD. Alternatively, DRN neuronal subpopulations are known to have distinct cortical and subcortical projections [11], and specific subpopulations targeted by viral injection may have affected which downstream regions contained tau spread.

Besides one study by Grinberg and colleagues[12], literature on AD neuropathology in the DRN of individuals without dementia or cognitive impairment is rare. The current study provides new evidence that, in addition to the LC, tau accumulates in the DRN at a premorbid stage and may be a prodromal indicator of dementia. The relatively low abundance of synuclein and the absence of TDP-43 in these individuals support the notion that the DRN may be more susceptible to tau aggregation, which is associated with AD and FTLD-tau. Accordingly, nearly all AD subjects had some degree of tau pathology in the DRN. We also demonstrate for the first time that tau pathology in the DRN drives depressive-like behaviors, serotonergic dysregulation, and astrocytic reactivity which may be precursors to neurodegeneration. Finally, tau was shown to propagate from the DRN to other subcortical regions and may act as a hub for the spread of tau pathology throughout the brain, a process that is accelerated by neural activity and astrocytic uptake of tau. Overall, these studies suggest that tau aggregation in the DRN may be an early indicator of AD, especially when combined with late-onset depression. Regular screening for late-onset depression may lead to an earlier diagnosis in individuals at risk for AD and could be used in concert with fMRI and PET imaging tools that can detect brainstem tau pathology or pathological changes in DRN integrity[70–72].

## Acknowledgements

We thank the Iowa Neurobank core for providing human post-mortem tissues for this study, Mariah Leidinger of the Comparative Pathology Core at The University of Iowa for providing histology services for human and mouse tissues, and the personnel of the Department of Pathology’s Immunohistochemistry Lab at The University of Iowa for performing TDP-43 staining. We additionally thank Ramasamy Thangavel for performing the α-syn-AT8-TH triple stain in human LC tissue.

## Declarations

### Ethics approval

All procedures on mice in this study were approved by the Institutional Care and Use Committee at the University of Iowa.

### Availability of data and materials

All data generated or analyzed during this study are included in this published article and its supplementary information files.

