## Supplemental Information for "Tau pathology in the dorsal raphe may be a prodromal indicator of Alzheimer’s disease"

**This PDF file includes:**

**Supplementary Methods**

**Supplementary Tables 1-3**

**Supplementary Figures and Legends S1-S4**

**References**

### Methods

#### *Animals*

Male and female C57BL/6J mice (Jackson Labs Stock 000664) were housed in University of Iowa animal facilities, under a 12-hour light-dark cycle with standard lighting and *ad libitum* access to food and water. Estrus cycle stages of female mice were tracked following every behavioral assay, using the method described by Byers et al. [1]. All procedures involving animals were in accordance with the University of Iowa's Institutional Animal Care and Use Committee.

#### *Stereotaxic surgery*

All surgeries were performed under aseptic conditions. At 8 weeks of age, mice were deeply anesthetized with 3% (v/v) isoflurane in oxygen and placed in a Leica Angle 2 stereotax (Germany) on a heating pad. Isoflurane was maintained at 1.5-2% thereafter. An adeno-associated viral vector (AAV) expressing the human transgene for P301L-tau or green fluorescent protein (GFP) was microinjected into the dorsal raphe nucleus (ML: 0.0; AP: 4.65; DV: -3.3 from Bregma at a 23.58º angle from the ML plane) using a 1 µL Hamilton syringe at 100nL/min (500 nL total). Post-operative analgesia consisted of two meloxicam injections (0.4 mg/kg, sc or ip) given 24 hours apart.

#### *Viral constructs*

The original plasmid used to construct the P301L-tau virus was a generous gift from Gloria Lee. Construction of the AAV (AAV2/8-CBA-P301Ltau-WPRE) was performed by the Viral Vector Core Facility at the University of Iowa. AAV2/8-CAG-GFP-WPRE was also obtained at the Iowa Viral Vector Core.

*Reverse transcriptase quantitative PCR (RT-qPCR)*

A separate cohort of C57BL/6J mice was injected with AAV expressing P301L-tau or GFP. Four weeks post injection, the mice were decapitated under isoflurane anesthesia and the brains were extracted and immediately flash frozen on dry ice. Micro-punches of the DRN were taken, and total RNA was isolated using TRIzol™ reagent (Thermo Fisher Scientific, MA, USA) as described previously [2]. DNA contamination was removed from the RNA using DNA-*free™* DNA removal Kit (Thermo Fisher Scientific). RNA purity and concentration were checked using a Nanodrop 1000 spectrophotometer (Thermo Fisher Scientific). Extracted RNA was reverse transcribed using the iScript cDNA Synthesis Kit (BioRad Laboratories, CA, USA) with the following thermal profile: 25°C for 5 minutes, 45°C for 20 minutes, and 95°C for 1 minute. SYBR green qPCR master mix (BioRad Laboratories) and specific primers for the target genes (Supplementary Table 1) were used for RT-qPCR. The following thermal profile was used on a CFX96 Real-time-PCR System (BioRad Laboratories): 10 minutes at 95°C; 40 cycles of 30 seconds at 95°C, 30 seconds at 60°C; and then followed with a melt curve analysis profile (60°C to 95°C in 0.5°C increments at a rate of 5 seconds/step). Fold changes in mRNA levels were determined by normalizing to β-actin Ct values using the $2^{-\Delta\Delta CT}$ method [3]. Samples below detection limit (determined by Ct values) were excluded from presented results. Results are shown as fold change in mRNA levels ± SEM.

*Behavior Assays*

All behavioral assays took place at least 4 weeks post-surgery and were performed in the following order (in males): Elevated Plus Maze, Open field, Social interaction, Sucrose Preference, Barnes Maze, Forced Swim, Contextual Fear condition. In females, all tests except the Barnes Maze and Forced Swim were performed. There was at least one day between subsequent behavior tests.

Elevated Plus Maze: Mice were placed in the center of an elevated, plus-shaped maze with open and enclosed arms (arms 35cm long, 5cm wide, 60cm off the ground; enclosed arms with 20cm high walls) and allowed to explore for five minutes as reported previously[4]. The assay was performed under dim light (~20 lux in open arms, and < 5 lux in closed arms) with the experimenter blocked from view using an opaque curtain. Sessions were recorded with a Basler GenICam and Media Recorder 4 software (Noldus, VA, USA). Time spent in open and closed arms and distance traveled were analyzed using EthoVision XT15 (Noldus).

Open Field test: Mice were placed in the corner of a white 50 x 50 x 25.5cm open box inside of a dimly lit (~ 20 lux), sound attenuated chamber and allowed to explore for 30 minutes[4]. Sessions were recorded with a Euresys piccolo U8H.264 camera mounted at the top of the sound attenuating chamber using Media recorder 4 software. The first 10 minutes of each session were analyzed for the following: time spent in the center of the box and total distance traveled.

Social Interaction test: Mice were placed in the center of a three-chambered, clear arena (64x41x21cm) inside of a large, dimly lit (~20 lux) sound-attenuated chamber. Mice acclimated to the arena and could explore all chambers for 10 minutes before a novel conspecific “stranger” mouse was placed under a small metal cage in the left or right chamber. Although the stranger was confined to one chamber under the cage, it was clearly visible to the test animal. An empty cage was placed in the opposite-side chamber [4, 5]. Mice were allowed to roam freely between all chambers for another 10 minutes and interact with the stranger or empty cage. Video was recorded using Media Recorder 4 software and an Euresys piccolo U8H.264 camera mounted at the top of the sound-attenuated chamber. Behavior was scored by measuring the time mice spent engaged in sniffing, circling, rearing, or otherwise in close proximity to and with their snouts pointed towards the stranger or empty cage.

Sucrose Preference test: Mice were placed in PhenoTypers (Noldus) equipped with lickometers and two bottles. One bottle was filled with distilled water and the other with 5% sucrose. The sucrose was kept on the same side each day. The bottoms of the PhenoTypers were lined with the same bedding as the home cages. The number of times the mice licked each bottle was recorded for 5 successive days, in 1-hour long sessions for 4 days, and a 20-minute test session on the 5th day. Lickometer data was acquired by Ethovision XT for each session. The first 10 minutes of this test session was used in the analysis.

Barnes Maze: A gray circular maze (91cm diameter, 93cm off the ground) had 20 holes evenly dispersed around the perimeter and was illuminated at ~580 lux. External to the maze were 3 visual cues: a yellow star on the screen that obstructed the experimenter from view, a blue square on the wall adjacent to the maze, and a green triangle on the wall behind the maze. On 3 consecutive days, mice were habituated to the maze and learned the location of a hidden compartment under one of the holes. On all training days, if a mouse did not enter the compartment on its own, it was gently nudged or placed inside by the experimenter. On the first day, mice were placed in the center of the arena with a clear beaker overtop for 30 seconds before being guided to the target and given 3 minutes to enter the chamber under the maze. A white noise was played until the mice entered the chamber and stopped upon entry, and they remained in the chamber for 1 minute. On the second day, each mouse received 3 trials of the following: 15 seconds in the center of the maze covered with an opaque box, followed by 2 minutes to freely move around the maze and enter the escape chamber. If the mouse did not enter the chamber, the clear beaker was placed overtop, keeping them within the vicinity of the chamber, and they were allowed 3 minutes to enter. Mice spent 1 minute in the chamber. While mice were exploring, a loud sustained 100 dB tone was played and then stopped upon chamber entry to serve as a mildly aversive stimulus to incentivize quick locating of and entry into the chamber. On the third day, mice received two of the previously described trials. Mice were given 1 day for consolidation and a break from training (Day 4). On the test day (Day 5), memory was evaluated. The escape chamber was removed, and animals were allowed to freely explore the arena for 2 minutes. Behavior was recorded by an overhead camera and acquired with Ethovision XT 11.5. The distance traveled, velocity, frequency of target and nontarget visits, and time spent in different quadrants of the maze were analyzed in Ethovision.

Forced Swim: The forced swim assay was used to evaluate depressive-like behaviors following injection of AAV-P301L-tau in the DRN of male mice. A tall, clear cylinder was filled with 24-25ºC tap water (32cm height x 20cm diameter; water height 25cm), and mice were gently placed inside for 6 minutes[4]. Following the test, mice were promptly removed from the cylinders and placed in a fresh, dry cage under a heat lamp for 5 minutes to allow them to warm up and dry off. Behavior was recorded from a side view camera and analyzed in Ethovision XT15. Six-minute videos were divided into two phases: a pretest (first 0-2 minutes), and test phase (next 2-6 minutes). Immobility was defined as the absence of movement (excluding tail flicks) for >1 s. Latency to first immobile bout, frequency of immobile bouts, and duration of bouts was measured in Ethovision.

Contextual Fear Conditioning*:* The fear conditioning assay was performed in sound-attenuated operant conditioning boxes manufactured by Ugo Basile (Model 46003, Italy) and recorded using a top-mounted infrared camera. A vanilla scent was added to the conditioning chamber as a contextual cue. On the conditioning day, mice were placed inside of the operant chamber and received a series of 5, 2-second long, 0.6 mA shocks at random intervals ranging from 1-2 minutes following a 2-minute baseline. Sessions ended 2 minutes after the final shock. Twenty-four hours later (recall or test day), mice were placed back in the same chamber for 10 minutes. Time spent freezing during the first 3, 5 and 10 minutes of this test session was determined by a trained observer blinded to the experimental condition. Freezing was scored at 5 s intervals and was defined by the absence of movement for at least 1 s. The data reported represent the first 3 minutes of the test session.

*Electrophysiology*

Brain Slice Preparation: Mice were deeply anesthetized with tribromoethanol (250 mg/kg, i.p.) and transcardially perfused with ice-cold, carbogenated modified artificial cerebrospinal fluid (aCSF) containing (in mM): 110 choline-Cl, 2.5 KCl, 7 $MgSO_4$, 0.5 $CaCl_2$, 1.25 $NaH_2PO_4$, 26.2 $NaHCO_3$, 25 glucose, 11.6 Na-ascorbate, 2 thiourea, and 3.1 Na-pyruvate (pH: 7.3 – 7.4; osmolality: 300 – 310 mOsmol/kg). Brains were quickly extracted and sliced with a vibratome (VT1200S; Leica Biosystems, Germany) to obtain coronal sections containing the DRN (thickness: 300 μm). The brain slices were recovered in the carbogenated choline-Cl-based aCSF (maintained at 34 °C) for 30 min. Afterwards, they were transferred to a different carbogen-saturated modified aCSF at room temperature in which they were held for at least 1 hour before recordings. The holding aCSF was made with (in mM): 92 NaCl, 2.5 KCl, 2 $MgSO_4$, 2 $CaCl_2$, 1.25 $NaH_2PO_4$, 30 $NaHCO_3$, 20 HEPES, 25 glucose, 5 Na-ascorbate, 2 thiourea, and 3 Na-pyruvate (pH: 7.3 – 7.4; 300 – 310 mOsmol/kg).

*Ex vivo* electrophysiological recordings: Neurons were visualized with a differential interference contrast (DIC) microscopy system (BX51W1; Olympus, Japan). Membrane currents were amplified by a Multiclamp 700 B amplifier (Molecular Devices, CA, USA), filtered at 3 kHz, and sampled at 20 kHz with a Digidata 1550B digitizer (Molecular Devices). Data were obtained using the pClamp 11 software (Molecular Devices). In the recording chamber, slices were continuously perfused (2 ml/min) with standard aCSF containing (in mM): 124 NaCl, 4 KCl, 1.2

$MgSO_4$, 2 $CaCl_2$, 1 $NaH_2PO_4$, 26 $NaHCO_3$, and 11 glucose (300 – 310 mOsmol/kg), saturated with 95% $O_2$/5% $CO_2$ and maintained at 30 ± 1°C.

In current clamp recordings for assessing serotonergic excitability, patch electrodes (3 – 5 MΩ) were filled with a solution made with (in mM): 135 K-gluconate, 5 NaCl, 2 $MgCl_2$, 10 HEPES, 0.6 EGTA, 4 $Na_2$-ATP, and 0.4 $Na_2$-GTP (pH: 7.3; ~290 mOsmol/kg). In some experiments, currents were injected into the neurons to hold the membrane potential at -70 mV. Input resistance was computed from the decrease in membrane potential upon hyperpolarizing current injection (-100 pA). Rheobase referred to the minimal current that could evoke an action potential. Evoked action potentials were tallied while injecting positive currents for 250 ms at 21 steps (0 – 200 pA, incremented by 10 pA). Data were reported as action potential frequency (number of spikes per 250 ms current step).

In voltage clamp recordings for examining synaptic transmission onto DRN 5-HT neurons, cesium methanesulfonate-based internal solution was used (in mM: 135 cesium methanesulfonate, 10 KCl, 1 $MgCl_2$, 10 HEPES, 0.2 EGTA, 4 Mg-ATP, 0.3 $Na_2$-GTP, 20 $Na_2$-phosphocreatine; pH 7.3, ~290 mOsmol/kg, with 1 mg/ml QX-314). Spontaneous excitatory postsynaptic currents (sEPSCs) were recorded continuously for 2 min while the neuron was voltage-clamped at -55 mV; similarly, in the same neuron, spontaneous inhibitory postsynaptic currents (sIPSCs) were recorded while the membrane potential was maintained at +10 mV.

To record electrically evoked EPSCs (eEPSCs) or IPSCs (eIPSCs), a glass pipette containing aCSF was placed lateral to the recorded neuron (distance: ~50 µm), and eEPSCs were evoked by square pulses (duration: 100 µs; 0.1 Hz) at different stimulating intensities (10, 20, 40, 80, 160, and 320 µA) while the neuron was voltage-clamped at -55 mV. Similarly, eIPSCs were evoked while the neuron was held at +10 mV. Moreover, to assess the probability of presynaptic neurotransmitter release, two stimuli with an inter-stimulus interval of 30 ms were delivered at 0.1 Hz, so eEPSCs and eIPSCs were evoked in pairs. The ratio of amplitudes of the two eEPSCs or two eIPSCs, i.e., paired pulse ratio (PPR = eEPSC2/eEPSC1 or eIPSC2/eIPSC1), was computed based on 10 – 30 consecutive trials. To identify 5-HT neurons *post hoc* by immunofluorescence, biocytin (2 mg/ml; Hello Bio, NJ, USA) was added into the internal solution, and only Tph2-positive neurons were used in data analysis.

*Histology*

Mouse Tissue Preparation: Mice were anesthetized with tribromoethanol (250 mg/kg, i.p.) and transcardially perfused with 30 ml 0.01 M phosphate buffered saline (PBS) followed by 30 ml 4% paraformaldehyde (PFA). Brains were extracted and post-fixed 24 hours in 4% PFA at 4°C, then transferred to PBS with 0.02% sodium azide. Tissues were paraffin-embedded and sectioned at 5 µm using a Leica rotary microtome at the University of Iowa Comparative Pathology Lab. For free-floating immunofluoescence, brains were transferred from 4% PFA to 15% and 30% sucrose solutions successively and allowed to equilibrate before embedding in OCT and sliced with a frozen sliding microtome (Leica).

Human Tissue

De-identified adult brain tissue was obtained from the University of Iowa NeuroBank or the NIH NeuroBioBank at the University of Maryland, Baltimore, MD. Since all samples were de-identified, this does not constitute human subjects research under the NIH Revised Common Rule. All cases had post-mortem intervals less than 24 hours. Alzheimer's disease and Lewy Body Dementia, as well as detailed staging information (Table 3 and Supplemental Table 1), were defined and determined according to standard criteria [6, 7]. All cases were reviewed by one of the authors (MMH), an experienced neurodegenerative and developmental neuropathologist.

DAB staining

All 3,3′-Diaminobenzidine (DAB) staining for ptau and α-synuclein in sectioned mouse and human tissue was performed by the University of Iowa Comparative Pathology Lab. TDP-43 DAB staining was carried out by the Department of Pathology's Immunohistochemistry Lab. DAKO autostainers (Agilent, CA, USA) were used by both labs, and all DAB-stained tissue was counterstained with hematoxylin after the DAB protocols.

*AT8 (phospho-tau Ser202/Thr205):* Paraffin-embedded slides were placed in distilled water ($dH_20$) and transferred to citrate buffer (pH 6.0) at 110°C for 15 minutes and allowed to cool for 20 minutes. After a 2-minute tap water rinse and then a 2-minute $dH_20$ rinse, slides were quenched with 3% hydrogen peroxide for 8 minutes and then rinsed again in distilled water. Next were 2 x 5-minute washes in 1X Dako Buffer, 20 minutes in Avidin, and 20 minutes in biotin with brief buffer rinses in between. For mouse tissue only, Rodent Block M (Biocare, CA, USA) was added for 20 minutes before 2 x 5-minute 1X Dako buffer washes. Slides were incubated in AT8 primary antibody (Table 2) diluted in Dako Diluent at 1:1000 for 15 minutes at room temperature (@RT) followed by 2 x 5-minute 1X Dako buffer washes, then mouse HRP secondary (EnVision, KS, USA) for 15 minutes. This was followed by 2 x 5-minute 1X Dako buffer washes, then 5 minutes in DAB, 3 rinses in 1X Dako Buffer, and finally a DAB enhancer for 3 minutes before a final wash in $dH_20$.

*TPH2*: Paraffin-embedded slides were placed in $dH_20$ and transferred to citrate buffer (pH 6.0) at 110°C for 15 minutes and allowed to cool 20 minutes. After a 2-minute tap water rinse and then a 2-minute $dH_20$, slides were washed in 1X Dako Buffer and quenched with 3% hydrogen peroxide for 8 minutes. After a brief $dH_20$ rinse, slides were washed twice in 1X Dako Buffer for 5 minutes. Next, slides were placed in Avidin and then biotin for 20 minutes each, with brief buffer rinses in between. Background Buster (Innovex Biosciences, CA, USA) was added to slides for 30 minutes, followed by 2x5-minute 1X Dako Buffer washes. Slides were incubated in TPH2 primary antibody (Table 2) diluted in Dako Diluent at 1:300 for 60 minutes @RT. After 2 x 5-minute 1X Dako buffer washes, rabbit HRP secondary (EnVision) was applied for 30 minutes. Following another 2 x 5-minute 1X Dako buffer wash, slides received DAB for 5 minutes, and then a rinse in 1X Dako buffer before a DAB enhancer was applied for 3 minutes. Slides received a final rinse in $dH_2O$.

*α-synuclein (pS129-α-synuclein):* The staining protocol for α-synuclein paraffin-embedded slides was the same as described for AT8 except for the blocking step. A 10% goat serum in 1X Dako buffer solution was used for 30 minutes of blocking. α-synuclein primary antibody (Table 2) was then diluted in Dako Diluent at 1:500 and applied to the slides for 15 minutes @RT. Rabbit HRP secondary (EnVision) was applied for 15 minutes.

*TDP-43 (pS409/410-TDP-43):* Staining was performed using the same protocol as AT8, with minor modifications. Quenching with peroxide was only 5 minutes. TDP-43 primary antibody (Proteintech, Germany) was diluted at 1:3000 and slides were incubated in this solution for 15 minutes @RT. Slides incubated in rabbit HRP secondary (EnVision) for 15 minutes. DAB was applied for 10 minutes.

Immunofluorescent staining

*Paraffin-embedded IF (IF-P):* The same staining protocol was used for mouse and human DRN paraffin-embedded IF staining, with slight alterations to primary antibody concentrations, choice of secondary antibodies, and secondary antibody concentrations (see Supplementary Table 2).

Slides were heated at 50-60°C to adhere tissue to slides and soften the paraffin. Paraffin was removed in two xylene washes. Tissue was then rehydrated starting with 100%, 90%, 70% ethanol and double distilled/Milli-Q water. Slides were placed in a water bath in polypropylene Coplin jars that contained antigen retrieval solution (0.514 g sodium citrate, 168 µL Tween 20, 200 mL MiliQ water) for 10 minutes. Slides were rinsed with 0.01 M PBS before hydrophobic barriers were drawn around tissue slices. Blocking solution (3% BSA, 0.4% Triton X-100 in PBS) was applied for one hour @RT and then slides were washed with PBS. Primary antibodies were diluted in the blocking solution and allowed to incubate overnight (18-24 hours) at 4°C in a humidified chamber at the following concentrations: AT8 1:1000 (mouse and human); TPH2 1:500

(both mouse and human); Iba1: 1:200; GFAP 1:1000. After three PBS washes, slides were incubated in secondary antibodies diluted at 1:500, 1:1000, or 1:2000 in blocking solution and protected from light for 1 hour @RT (See Suppl. Table 2 for full antibody information). Slides were once again washed with PBS and then coverslipped with Vectashield Antifade mounting medium (Vector Labs, CA, USA) and sealed with clear nail polish.

*Human LC IF-P:* The paraffin sections were deparaffinized with xylene and rehydrated with series of changes in ethanol. Then sections were subjected to heat-induced antigen retrieval in sodium citrate buffer solution for 10 min. Sections were incubated in blocking solution of 2% donkey serum+0.3% Triton-X 100 in PBS for 1h at room temperature. Primary antibodies were used to localize tyrosine hydroxylase positive neurons with anti-Sheep TH antibody [1:500] in combination with phosphorylated tau with AT8 antibody [1:1000] to identify the neurofibrillary tangles, and phosphorylated forms of alpha-Synuclein antibody [1:1000] to detect the Lewy bodies in the LC area (See Suppl Table 2 for full antibody details). Sections were incubated for overnight at 4°C with primary antibodies mixture. After washing with PBS, sections were incubated for 1h @RT in the dark with cocktail mixture of secondary antibodies of Alexa Fluor 488 donkey anti-sheep [1:500] for anti-TH, Alexa Fluor 568 donkey anti-mouse [1:500] for anti-ptau and Alexa Fluor 647 donkey anti-rabbit [1:1000] α-syn. After thorough washing with PBS, sections were cover slipped with prolong diamond antifade mounting medium with DAPI (P36962, Invitrogen). Images were taken using a confocal microscope (Zeiss) with identical settings for each image (40X objective, 1.7X zoom).

*Free-floating IF (IF-F):*

*AT8-NeuN-GFAP-Iba1:* 35μm DRN tissue slices were rinsed in PBS 3 times for 5 mins, then permeabilized using 0.5% Triton X-100 in PBS. Tissue was rinsed 2X 5min in PBS before blocking for 1 hour @RT. (Blocking buffer: 10% normal donkey serum (Jackson ImmunoResearch), 2% FcX receptor blocker (BioLegend), 0.1% Trion X-100 in PBS). Primary antibodies were diluted in incubation buffer (10% normal donkey serum (Jackson ImmunoResearch), 0.1% Trion X-100 in PBS) as listed in Table 2 and incubated overnight @4°C [AT8: 1:1000; NeuN: 1:2000 ;GFAP: 1:1000; Iba1: 1:200]. Slices were then washed 3X 5 min in PBS, followed by secondary antibody incubation for 2 hour @RT [diluted in PBS 1:500]. Finally, tissue was washed 4X 10 mins in PBS prior to mounting on slides and cover slipped using Vectashield Vibrance (Vector Labs).

*HT7-SERT*: 35µm tissue slices spanning regions downstream of the DRN were permeabilized and blocked as described above. Primary antibodies were diluted in incubation buffer (same as above) for 3 overnights @4°C [HT7: 1:1000; SERT: 1:2500]. Slices were then washed 4 X 5 min in PBS, followed by secondary antibody incubation for 2 hour @RT [diluted in PBS 1:500]. Subsequent washes and tissue mounting was the same as described above.

*Confocal microscopy*

All fluorescent images of human DRN and mouse tissue were taken on an Olympus Fluoview FV3000 confocal microscope using Olympus objective lenses (UPLSAPO10X2, NA 0.4; UPLXAPO20X, NA 0.8; UPlanXApo40X, NA 1.4). AT8-TPH2 z-stack images for analysis were taken at 10X, 1.5 optical zoom, step size 1.0 µm. AT8-GFAP-Iba1 z-stack images for analysis were taken at 20X, step size 0.5 µm. Human tissue IF z-stack images were taken at 20X, 2X optical zoom, step size 1.0 µm. Mouse 35µm AT8-NeuN-GFAP-Iba1 and HT7-SERT z-stack images were taken at 40X, 1.0 µm step size. Human DAB images were taken on an Olympus brightfield microscope.

*Image Processing and Analysis*

Preprocessing of images for analysis was performed in ImageJ. All images were adjusted to the same brightness and contrast settings and subjected to consistent bright outlier removal and de-speckling as necessary.

Optical density: Image analysis of paraffin-embedded tissue was done in ImageJ (NIH). Optical Density (OD) values were obtained following the standard protocol found on the ImageJ NIH website, with OD of the image background subtracted to generate a "Net OD".

Semi-quantitation of DAB images: Images were scored blinded with supervision and assistance from a certified neuropathologist (MMH). Tau pathology in mouse tissue was designated as follows: Multiple areas in multiple fields of view (++); Very little pathology localized to single fields of view (+); No pathology (-). Human tau pathology was additionally semi-quantitatively scored by an experienced, blinded neuropathologist (MMH) (Table 2): 0: none ;1: rare ; 2: frequent. Control cases 4 and 6 exhibited extremely rare pathology, and were assigned a score above 0.

TPH2-AT8 analyses: TPH2+ immunostaining defined the DRN for analysis of AT8 and TPH2 in mouse tissue, and the injection site (GFP or AT8 immunostaining) defined regions for analyses of immunological markers. Representative images for figures were prepared in Adobe Photoshop with adjustments to image orientation (Straightening), Hue/Saturation, Brightness/Contrast, and Vibrance.

AT8-GFAP-Iba1-NeuN: Analysis of AT8+ staining that was also positive for one of 3 cell-type markers (GFAP, Iba1, NeuN) was done in QuPath [8], followed by percent of total calculations in excel: $\%\,Positive = \frac{\#\,objects\ co{-}positive\ for\ AT8{+}Marker}{total\ \#\ AT8{+}objects} * 100$. Representative images were adjusted for Brightness/Contrast in ImageJ.

HT7-SERT: Total area of HT7 and SERT immunofluorescence was determined by pixel threshold in QuPath, followed by the total area of SERT immunofluorescence within the HT7 area. $\%\ \text{HT7 that is SERT+} = \frac{SERT\ area\ within\ HT7}{total\ HT7{+}area} * 100$

*Statistical Analysis:* For comparisons between two groups with one independent variable, Student t-tests were used where appropriate; otherwise Mann-Whitney tests or Kolmogorov-

Smirnov (K-S) tests were used. ANOVA was used for all analyses involving two or more independent variables. Bonferroni tests were used for post hoc pairwise comparisons. Statistical significance was ascertained when $p<0.05$. Statistical significance was defined as $^*p<0.05$. $^{**}p<0.01$, $^{***}p<0.001$, $^{****}p<0.0001$. Data are expressed as mean ± SEM unless otherwise specified.

### Supplementary Tables

### Supplementary Table 1: RT-qPCR primers

| Gene Name | Forward/Reverse (5'-3') | Sequence |
|---|---|---|
| *β-actin* | Forward | CCAGCCTTCCTTCTTGGGTA |
| | Reverse | GAGGTCTTTACGGATGTCAACG |
| *Il1a* | Forward | TTGCTGAAGGAGTTGCCAGA |
| | Reverse | GCACCCGACTTTGTTCTTTGG |
| *Il1b* | Forward | GCCACCTTTTGACAGTGATGAG |
| | Reverse | AAGGTCCACGGGAAAGACAC |
| *Il6* | Forward | GAGACTTCCATCCAGTTGCCT |
| | Reverse | TCCTCTGTGAAGTCTCCTCTCC |
| *Il12a* | Forward | ATCACACGGGACCAAACCAG |
| | Reverse | CCAAGGCACAGGGTCATCAT |
| *Il12b* | Forward | TCATCAAACCAGACCCGCC |
| | Reverse | TCTGGTTACACCCCTCCTCT |
| *Ifn-γ* | Forward | TTGCGGGGTTGTATCTGGG |
| | Reverse | TGGCCCGGAGTGTAGACAT |
| *Tnf-α* | Forward | CGGGCAGGTCTACTTTGGAG |
| | Reverse | ACCCTGAGCCATAATCCCCT |
| *Nfκb1* | Forward | ACTGTCTGCCTCTCTCGTCT |
| | Reverse | CCGTGGGGCATTTTGTTCAG |
| *Nfκb2* | Forward | TCCCGAATGGACAAGACAGC |
| | Reverse | GGAGAGAAGTCCCCAAAGGC |
| *Tph2* | Forward | GACCCAAAGACGACCTGCTT |
| | Reverse | CTGCGTGTAGGGGTTGAAGT |
| *Frk* | Forward | AGCAGGTCAGGAAGAAGCAC |
| | Reverse | CTCACCATACCTCCCGCTTC |

**Supplementary Table 2: Antibody Information**

| Stain | Tissue Type | Antibody | Concentration |
|---|---|---|---|
| **AT8-TPH2 IF-P** | Mouse | Mouse anti-phosphotau, Thermofisher MN1020 | 1:1000 |
| | | Cy3 Donkey Anti-Mouse<br>Jackson ImmunoResearch 715-165-150 | 1:500 |
| | | Rabbit anti-tryptophan hydroxylase<br>Novus NB100-74555 | 1:500 |
| | | DyLight™ 405 AffiniPure Donkey Anti-Rabbit<br>Jackson ImmunoResearch 711-476-152 | 1:500 |
| **AT8-Iba1-GFAP IF-P** | Mouse | Mouse anti-phosphotau, Thermofisher MN1020 | 1:1000 |
| | | DyLight™ 405 AffiniPure Donkey Anti-Mouse<br>Jackson ImmunoResearch 715-475-150 | 1:2000 |
| | | Rabbit anti-Iba1, Abcam ab178846 | 1:200 |
| | | Cy3 Donkey Anti-Rabbit<br>Jackson ImmunoResearch 711-165-152 | 1:500 |
| | | Chicken anti-GFAP, Abcam ab4674 | 1:1000 |
| | | Alexa Fluor 647 Donkey Anti-Chicken<br>Jackson ImmunoResearch 703-605-155 | 1:2000 |
| **AT8-NeuN-GFAP-Iba1 IF-F** | Mouse | Mouse anti-phosphotau, Thermofisher MN1020 | 1:1000 |
| | | Donkey Anti-Mouse Alexa Fluor647<br>Abcam ab150107 | 1:500 |
| | | Guinea Pig anti-NeuN, Millipore ABN90 | 1:2000 |
| | | Alexa Fluor 488 Donkey Anti-Guinea Pig<br>Jackson ImmunoResearch 706-545-148 | 1:500 |
| | | Chicken anti-GFAP, Abcam ab4674 | 1:000 |
| | | DyLight™ 405 AffiniPure Donkey Anti-Chicken<br>Jackson ImmunoResearch 703-475-155 | 1:500 |
| | | Rabbit anti-Iba1, Abcam ab178846 | 1:200 |
| | | Donkey anti-Rabbit Alexa Fluor 555<br>Invitrogen A31572 | 1:500 |
| **AT8-SERT IF-F** | Mouse | Mouse anti-Tau monoclonal antibody (HT7),<br>Invitrogen MN1000 | 1:1000 |
| | | Alexa Fluor 488 AffiniPure Donkey Anti-Mouse<br>Jackson ImmunoResearch 715-545-150 | 1:500 |
| | | Rabbit anti-serotonin transporter<br>Immunostar 24330 | 1:2500 |
| | | Donkey anti-Rabbit Alexa Fluor 555<br>Invitrogen A31572 | 1:500 |

**Supplementary Table 2: Antibody Information continued.**

| Stain | Tissue Type | Antibody | Concentration |
|---|---|---|---|
| **AT8-TPH2 IF-P** | Human DRN | Mouse anti-phosphotau, Thermofisher MN1020 | 1:1000 |
| | | Alexa Fluor 555 Goat Anti-Mouse, Abcam Ab150114 | 1:1000 |
| | | Rabbit anti-tryptophan hydroxylase Novus NB100-74555 | 1:500 |
| | | Alexa Fluor 488 Goat Anti-Rabbit, Abcam Ab150077 | 1:1000 |
| **α-syn-AT8-TH IF-P** | Human LC | Rabbit anti-Alpha Synuclein (phospho-S129) Abcam ab51253 | 1:1000 |
| | | Alexa Fluor 647 Donkey anti-Rabbit Invitrogen A31573 | 1:1000 |
| | | Mouse anti-phosphotau, Thermofisher MN1020 | 1:1000 |
| | | Alexa Fluor 568- Donkey anti-mouse Invitrogen A10037 | 1:500 |
| | | Sheep anti-Tyrosine Hydroxylase Sigma-Millipore- AB1542 | 1:500 |
| | | Alexa Fluor 488 Donkey anti-Sheep Invitrogen A11015 | 1:500 |
| **AT8 DAB** | Mouse & Human (DRN&LC) | Mouse anti-phosphotau, Thermofisher MN1020 | 1:1000 |
| | | HRP Mouse, EnVision K4001 | NA |
| **TPH2 DAB** | Human DRN | Rabbit anti-tryptophan hydroxylase Novus NB100-74555 | 1:300 |
| | | HRP Rabbit, EnVision K4003 | NA |
| **α-synuclein DAB** | Human DRN&LC | Rabbit anti-Alpha Synuclein (phospho-S129) Abcam ab51253 | 1:500 |
| | | HRP Rabbit, EnVision K4003 | NA |
| **TDP-43 DAB** | Human DRN&LC | Rabbit anti-Phospho-TDP-43 (Ser409/410) Proteintech 22309-1-AP | 1:3000 |
| | | HRP Rabbit, EnVision K4003 | NA |

IF-P = Immunofluorescence, paraffin-embedded

IF-F = Immunofluorescence, free-floating

**Supplementary Table 3: Full list of regions with observed human tau spread**

| 4 weeks post virus injection | | 8 weeks post virus injection | |
|---|---|---|---|
| AHiPM | PH | AHC | P1Rt |
| BLA | PIF | AHP | PaF |
| BMP | PIL | AIP | PAG |
| CeA | Pir | Arc | PaMP |
| Cg | Pn | BLA | PBP |
| CM | Po | CA1 | PG |
| CPu | PoMn | CA3 | PH |
| DS/post/VS | PP | CeA | PIF |
| Ect | PrGMC | Cg | Pir |
| Ent | PRh | CM | PMV |
| GP | PV | cp | PN |
| ic | PVP | CPu | poDG |
| LH | REn | DLG | PP |
| LHb | rmx | DM | PrEW |
| MeA | S1ULp/S2 | DMC | PrGMC |
| mfb | SG | DMV | PSTh |
| MGN | SN | DpG | PV |
| MHb | TeA | DS/post/ VS | PVG |
| mRt | VPM | DTM | Re |
| Or | ZIC | ec | Rh |
| P1PAG | ZID | Ent | RLi |
| PaF | ZIR | fi | RMC |
| PAG | ZIV | IMD | RPC |
| PBP | | LaDL | SN |
| | | LD | st |
| | | LH | SubI |
| | | LPAG | SuG |
| | | MeA | TeA |
| | | MGN | VTA |
| | | mlf | ZIC |
| | | MoDG | ZID |
| | | mRt | ZIV |
| | | Or | |

Region abbreviations based on Paxinos and Frankin's *The Mouse Brain*, Fifth Ed.

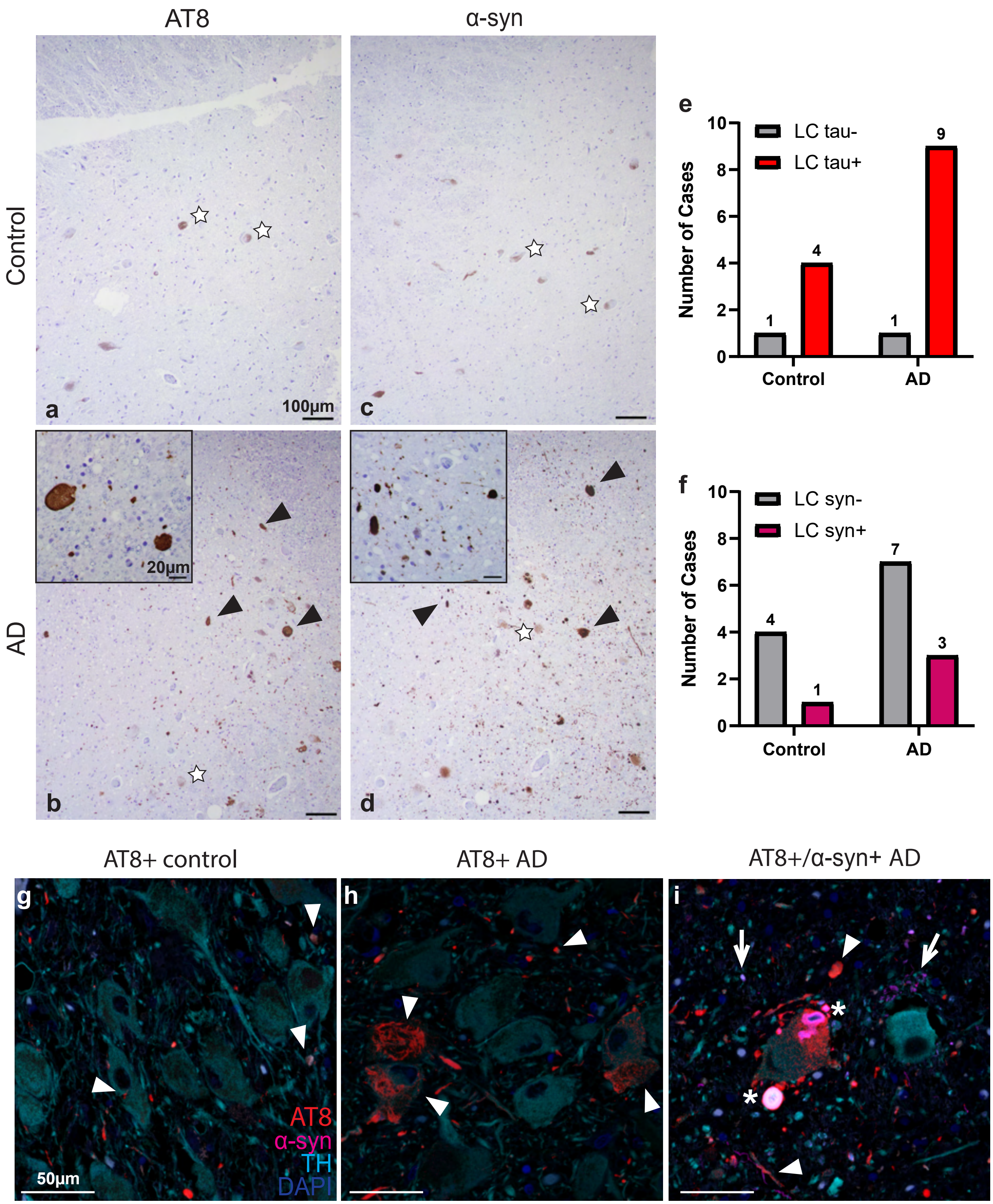
AT8
α-syn
Control
AD
a
b
c
d
100µm
20µm
e
Number of Cases
LC tau-
LC tau+
Control
AD
f
LC syn-
LC syn+
AT8+ control
AT8+ AD
AT8+/α-syn+ AD
g
h
i
AT8
α-syn
TH
DAPI
50µm

**Suppl. Fig. 1 Neurodegenerative markers in the locus coeruleus of control and AD patients.** DAB immunohistochemistry for ptau (AT8; a,b) and α-synuclein pathology (α-syn; c,d) in one control (a,c) and AD patient (b,d). Stars indicate DAB+ neuromelanin staining, characteristic of noradrenergic cells of the LC. Black arrowheads denote DAB+ pathology for AT8 or α-syn. Counter stain is hematoxylin. Large images were taken at 10X and scale bars are 100μm. Inset scalebars are 20μm and images were taken at 40X. **e** Bar graph showing the number of control and AD cases positive for AT8 in the LC. 4/5 control cases and 9/10 AD cases exhibited AT8 pathology in the LC. **f** Bar graph showing the number of control and AD cases positive for α-syn in the LC. Only 1/5 control cases and 3/10 AD cases had α-syn pathology in the LC.TDP-43 images are not pictured, as all cases were negative for staining in the LC. **g-i** Immunofluorescent stain of AT8, α-syn, and TH in human LC. **g** Control case positive for AT8 in the LC. **h** AD case positive for AT8 in the LC. **i** AD case positive for AT8 and α-syn in the LC. White arrowheads point to ptau and TH double-labeling. White arrows indicate α-syn and TH double-labeling.  Asterisks indicate triple labeling of ptau, α-syn, and TH. Images taken at 40X, 1.7 zoom. Scalebar=50 μm. red=AT8, magenta=α-syn, cyan=TH, blue=DAPI

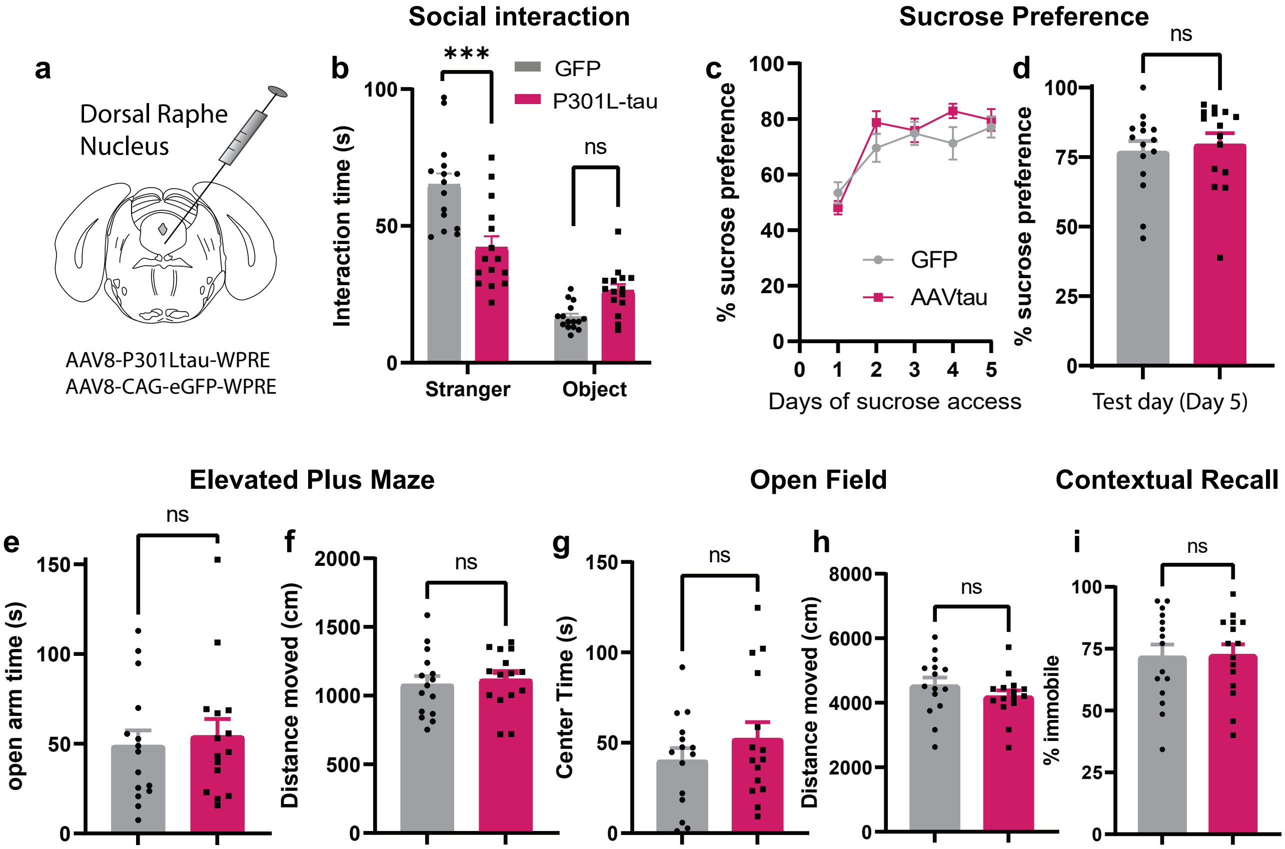


**Suppl. Fig. 2 DRN tau pathology leads to social deficits in female mice.** Female C57BL/6J mice received infusion of the same P301L-tau or GFP virus utilized in male mice (n=15 per group, 30 total). **a** Schematic of viral infusion into the DRN of female mice. **b** Time female mice spent interacting with a stranger mouse and object in the social interaction assay. **c** X-Y plot of sucrose preference across all days of the sucrose preference assay. **d** Bar graph of percent sucrose preference on the final day of testing (day 5). **e-f** Bar graphs displaying time spent in the open arms (e) and distance moved (f) in the EPM. **g-h** Bar graphs of the time spent in the center of the arena (g) and distance moved (h) in the OFT. **i** Bar graph showing the percent time spent immobile in the contextual fear conditioning task. Data is represented as mean ± SEM. ***$p<0.001$.

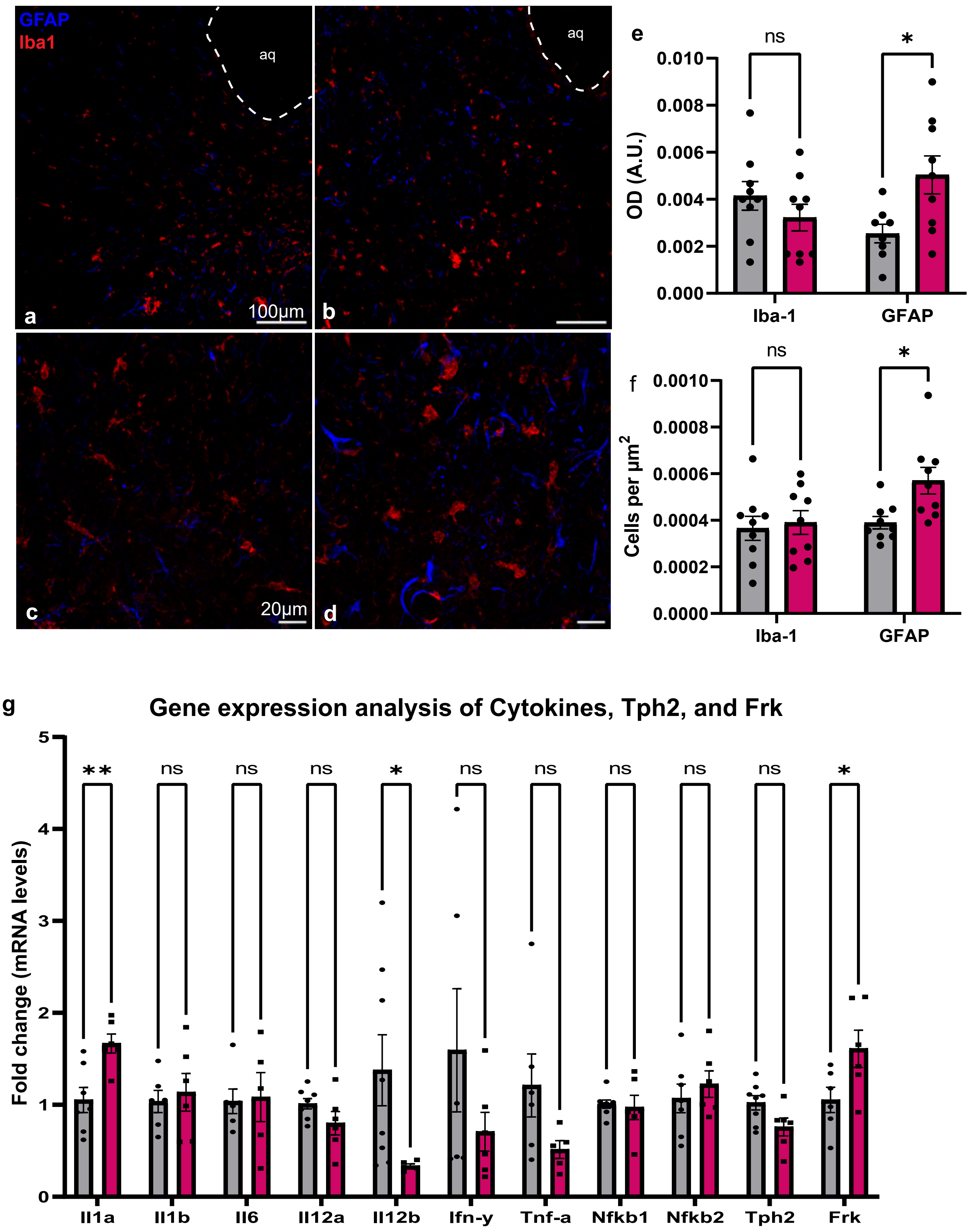
GFAP
Iba1
aq
aq
100µm
20µm
a
b
c
d
e
ns
*
OD (A.U.)
0.010
0.008
0.006
0.004
0.002
0.000
Iba-1
GFAP
f
ns
*
Cells per µm2
0.0010
0.0008
0.0006
0.0004
0.0002
0.0000
Iba-1
GFAP
g
Gene expression analysis of Cytokines, Tph2, and Frk
Fold change (mRNA levels)
5
4
3
2
1
0
**
ns
ns
ns
*
ns
ns
ns
ns
ns
*
Il1a
Il1b
Il6
Il12a
Il12b
Ifn-y
Tnf-a
Nfkb1
Nfkb2
Tph2
Frk

**Suppl. Fig. 3 Astrocytic activation and neuroinflammation in the DRN of P301L-tau$^{DRN}$ mice. a-b** Immunofluorescent staining for astrocytic marker GFAP (blue) and microglial marker Iba1 (red) in GFP (a) and P301L-tau animals (b) at 20X magnification. **c-d** GFAP and Iba1 IF in GFP (c) and P301L-tau mice (d) at 40X magnification. **e** Bar graphs displaying mean OD of GFAP+ and Iba1+ staining in ROIs expressing AT8. **f** Bar graph displaying GFAP+ and Iba1+ cell density in ROIs. Data are represented as mean ± SEM. Scalebar in a,b= 100 µm; c,d = 20 µm. aq=cerebral aqueduct. *$p<0.05$. **g** Gene expression analysis of cytokines, Tph2 and frk in the DRN of GFP and P301L-tau injected mice.

**a**

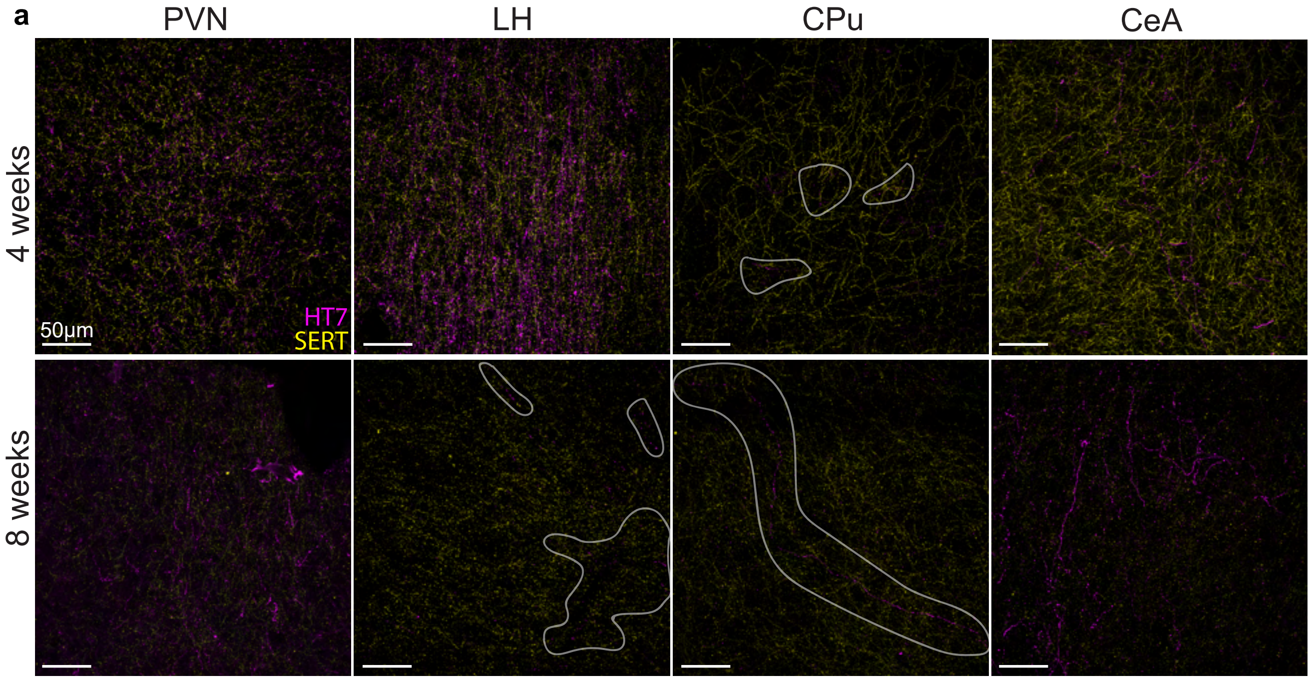


**b**

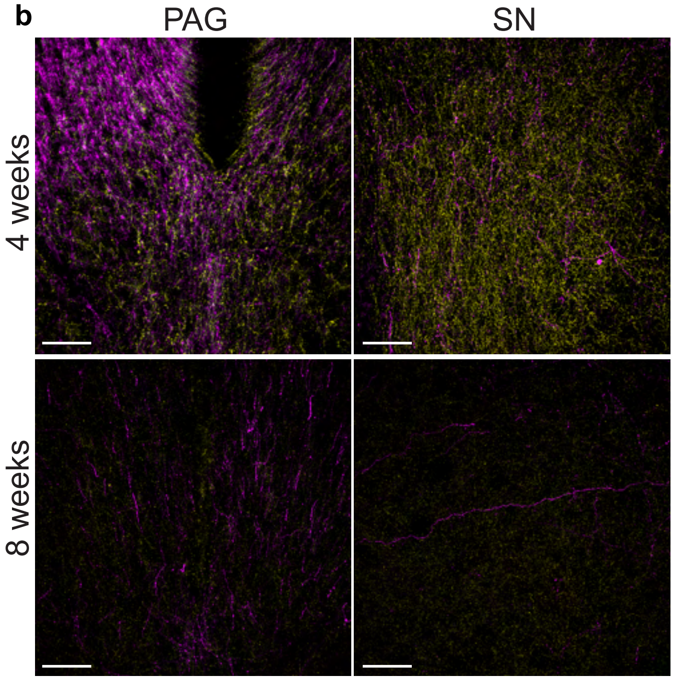


**c** (4 weeks)

| | HT7 % IR | % HT7 that's SERT+ | SERT % IR | % SERT that's HT7+ |
|---|---|---|---|---|
| HP | 0.00 | 0.00 | 12.35 | 0.00 |
| Thal | 11.47 | 25.63 | 22.11 | 13.21 |
| HT | 9.01 | 29.22 | 25.69 | 10.04 |
| SN | 6.44 | 51.26 | 47.46 | 7.02 |
| PAG | 38.90 | 22.58 | 19.10 | 43.33 |
| CPu | 0.08 | 19.88 | 12.71 | 0.11 |
| Amy | 3.15 | 16.20 | 14.71 | 3.26 |
| Ctx | 0.41 | 31.00 | 24.07 | 0.63 |

**d** (8 weeks)

| | HT7 % IR | % HT7 that's SERT+ | SERT % IR | % SERT that's HT7+ |
|---|---|---|---|---|
| HP | 0.38 | 6.91 | 4.40 | 0.60 |
| Thal | 1.39 | 12.37 | 12.74 | 1.41 |
| HT | 0.25 | 22.36 | 23.65 | 0.23 |
| SN | 2.01 | 4.51 | 4.72 | 1.94 |
| PAG | 4.63 | 6.26 | 5.50 | 5.28 |
| CPu | 0.53 | 9.54 | 7.43 | 0.64 |
| Amy | 1.38 | 31.78 | 24.38 | 1.57 |
| Ctx | 0.00 | 0.00 | 6.05 | 0.00 |

**Suppl. Fig. 4 P301L-tau$^{DRN}$ co-labels with SERT in DRN-connected regions. a-b** Immunofluorescent staining for human tau (HT7, magenta) and serotonin reuptake transporter (SERT, yellow) in regions downstream of the DRN in P301L-tau$^{DRN}$ mice 4 weeks (top rows) and 8 weeks (bottom rows) post-viral injection. In the CPu and LH, translucent outlines are included to indicate HT7+ staining that may be difficult to identify at 100% figure scale. Images taken at 40X. Scalebar=50 µm. **c** Table containing percent immunoreactive area (% IR) of human tau (HT7) and SERT in brain regions after 4 weeks of viral transduction, as well as the percent of total HT7+ or SERT+ area that is co-labeled with the other. **d** Table containing the same information as c, but for brain regions 8 weeks post-injection. A comprehensive list of regions where tau was observed to spread is included in Suppl Table 3. Abbreviations: PVN=paraventricular nucleus; LH=lateral hypothalamus; CPu=caudate/ putamen; CeA=central nucleus of the amygdala; PAG=periaqueductal gray; SN=substantia nigra; HP=hippocampus; Thal=thalamus; HT=hypothalamus; AMY=amygdala; Ctx=cortex
